# The evolution of autonomy from two cooperative specialists in fluctuating environments

**DOI:** 10.1101/2023.08.24.554579

**Authors:** Xiaoli Chen, Miaoxiao Wang, Laipeng Luo, Xiaonan Liu, Liyun An, Yong Nie, Xiao-Lei Wu

## Abstract

From microbes to humans, organisms perform numerous tasks for their survival, including food acquisition, migration, and reproduction. A complex biological task can be performed by either an autonomous organism or by cooperation among several specialized organisms. However, it remains unclear how autonomy and cooperation evolutionarily switch. For example, it remains unclear whether and how a group of cooperative specialists can evolve into an autonomous organism. Here, we address this question by experimentally evolving a mutualistic microbial consortium composed of two specialists that cooperatively degrade naphthalene. We observed that autonomous genotypes capable of performing the entire naphthalene degradation pathway evolved from two cooperative specialists and came to dominate the microbial community. This evolutionary transition was driven by the horizontal gene transfer between the two specialists. However, this evolution was exclusively observed to occur in the fluctuating environment supplied with either naphthalene or pyruvate, where mutualism and competition between the two specialists alternated. The naphthalene-supplied environment exerts selective pressure that favors the expansion of autonomous genotypes but was associated with both low cell density and low coexistence levels. In contrast, the pyruvate-supplying environment promoted the coexistence and cell density of the cooperative specialists, thereby increasing the likelihood of horizontal gene transfer. Using a mathematical model, we quantitatively demonstrate that environmental fluctuations facilitate the evolution of autonomy through HGT when the relative growth rate and carrying capacity of the cooperative specialists allow enhanced coexistence and higher cell density in the competitive environment. Together, our results demonstrate that mutualistic cooperation can evolve into autonomous organisms through direct genetic exchange under specific conditions, including the alternation of mutualism and competition in fluctuating environments in a manner frequently occurring in nature.

## Introduction

Bio-organisms perform diverse biological tasks in response to numerous pressures for survival, from hunting for food to reproducing and raising offspring. Biological tasks can be autonomously completed by one organism itself, or through the cooperation of social organisms working together for common or mutual benefits^1,2^. It is commonly observed that a specific task is achieved by either of the two strategies in different organisms. For example, many animal predators, such as tigers (*Panthera tigris*)^3^, cheetahs (*Acinonyx jubatus*)^4^, and golden eagles (*Aquila chrysaetos*)^5^, are thought to hunt alone. In contrast, predators like lions (*Panthera leo*)^6^, hyenas (*Hyaenidae*)^7^, killer whales (*Orcinus orca*)^8^, and Harris’ Hawks (*Parabuteo unicinctus*)^9^, prefer to form parties composed of multiple individuals to hunt via cooperation. Bacteria operate the nitrification pathway to acquire available energy, which is often conducted by mutualistic cooperation (refers specifically to the cooperation between different populations): ammonia-oxidizing bacteria convert ammonia to nitrite and nitrite-oxidizing bacteria convert nitrite to nitrate^10^. However, *Nitrospira* bacteria was shown to achieve complete nitrification within a single cell^11-13^. Similarly, anaerobic degradation of long-chain alkanes can be either performed via mutualistic cooperation between alkane-oxidizing bacteria and methanogenic archaea (known as syntrophy)^14^ or by an autonomous archaeon, namely *Candidatus* Methanoliparum^15^.

Previous studies suggested that the survival strategy of organisms, whether individual autonomy or cooperative interaction, is determined by the environmental conditions in which they reside, with the aim of enhancing their overall success in accomplishing specific tasks^16,17^. For instance, bacteria autonomously performing nitrification are favored in biofilms because biofilms restrain the spatial intermixing of cooperative specialists, and diffusion barriers in biofilms limit the exchange of metabolites among different individuals, which is necessary for metabolic cooperation^13,18^. However, whether and how autonomy and cooperation switch on evolutionary time scales remains to be elucidated. For example, when cooperation fosters higher productivity, an autonomous organism can evolve into different cooperative specialists. Reversely, when an autonomous organism is selectively favored by specific environmental conditions, can the group of cooperative specialists evolve into the autonomous organism? Such evolutionary transitions are essential for bio-organisms to adapt to their changing habitats. Several hypotheses have been proposed to explain how cooperative groups evolved from an autonomous ancestor. For example, the Black Queen hypothesis predicts that the evolution of cooperation within microbial communities is originally driven by the leakiness of public goods ^19,20^. This prediction explains the evolution of cooperation in several cases^21-24^ and has been tested by a few conceptual models^17,19,25^, as well as in the experimental evolution of *Pseudomonas aeruginosa*^26^ and engineered *Saccharomyces Cerevisiae*^27^.

In comparison, our understanding of how cooperative specialists evolve into autonomously functioning organisms remains inadequate. At present, two distinct pathways are likely to be involved in this process^28^. In the first pathway, one of the cooperative specialists coopts its missing function by evolving this function itself. One study observed this pathway in the experimental evolution of pellicle biofilms of *Bacillus subtilis*, in which one of the ancestral cooperative genotypes evolved to form biofilm autonomously^29^. In the alternative path, one cooperative specialist obtains the lacking function from its cooperative partner, for example, by horizontal gene transfer (HGT)^30,31^. Genomic studies of microorganisms suggested that mobile genetic elements (plasmids, bacteriophages, transposons) transmitted via HGT often carry multiple traits that complement the functional deficits of the auxotrophic organisms^32-34^. One recent study assessed the effects of HGT on bacterial metabolic networks using 835 genomes^35^. That study revealed that HGT allows bacteria to add new catabolic routes to bacterial metabolic networks, potentially augmenting the function of the bacteria. These genomic analyses strongly suggest that autonomy can evolve from mutualistic cooperation, namely through genetic transfer between different organisms. However, clear experimental evidence supporting this hypothesis remains absent.

Another crucial aspect to consider is the impact of environmental fluctuations that frequently occur in natural habitats^36^. These fluctuations have been shown to influence the co-evolution of different genotypes^37^. First, the fitness of a novel mutant is varied among different environments. In a stable environment, the fate of mutation is mostly decided by its fitness in that environment. In comparison, whether a mutation can be fixed in a fluctuating environment is determined by both its average fitness across environments and the frequency of environmental fluctuations^38^. Second, environmental fluctuations can also influence the coexistence of different genotypes, because relative fitness and ecological interactions between different genotypes can vary in different environments^36^. Several recent studies suggested that environmental fluctuations promote the coexistence of different genotypes^39-41^. In such cases, environmental fluctuations may benefit the coevolution of different genotypes, for example, by improving the HGT rate among these genotypes. The evolution of autonomy from cooperation requires that: 1) the evolved autonomous genotypes possess higher overall fitness (adaption); 2) the cooperative specialists can co-exist to enable HGT. Environmental fluctuations can influence both of these aspects.

Here, we set out to experimentally test whether a mutually cooperative consortium can evolve into an autonomous organism and how environmental fluctuations influence the evolution. We performed this study by evolving a mutualistic consortium composed of two bacterial specialists that cooperatively degrade naphthalene in constant and fluctuating environments. Based on our experimental findings, we built a mathematical model to search the key factors that determine whether autonomy can evolve from mutualistic cooperation.

## Results

### Design of a synthetic consortium and the experimental evolution

To investigate whether a mutually cooperative consortium can evolve into an autonomous organism, we made modifications to a synthetic mutualistic consortium that was originally employed in our prior experimental study^42^. In this consortium, the first strain (referred to as Degrader thereafter) degrades naphthalene to salicylate, which is further decomposed into small molecules such as pyruvate by the second strain. Degrader is able to grow alone using naphthalene as the sole carbon source while the second strain cannot (Figure 1A-B; Figure S1). Notably, we found that salicylate is toxic to both strains^43,44^ (Figure S2), thus indicating that the second strain also contributes to the detoxification process (hence we refer to the second strain as Detoxifier thereafter; Figure 1B), as confirmed by our coculture experiment (Figure S2). Similar forms of cooperative degradation of organic compounds have been frequently observed^45-53^. These results suggested that the two strains showed facultative mutualism upon co-culture (Figure S1). We engineered the two strains in this consortium from an ancestral *Pseudomonas stutzeri* strain, which can completely degrade naphthalene into small molecules (referred to as Autonomist thereafter; Figure 1C). When culturing Autonomist or the mutualistic consortium using naphthalene as the sole carbon source, Autonomist exhibits a higher growth rate and biomass yield than those of the consortium (Figure 1A).

**Figure 1.**
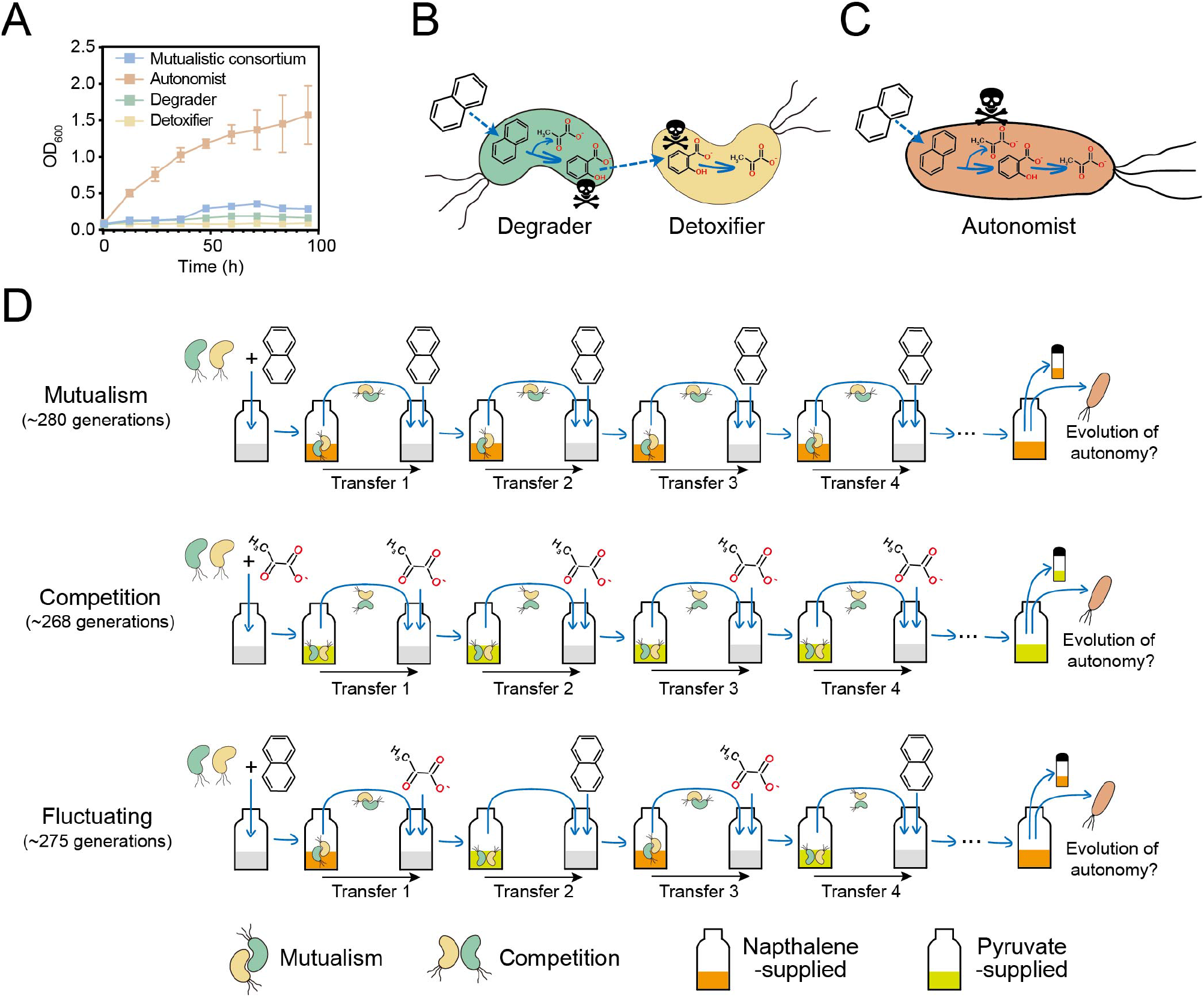
Experimental design. (A) The growth curve of the synthetic consortium and the Autonomist strain with naphthalene as the sole carbon source. (B) Schematic diagram of our synthetic consortium performing mutualistic cooperation. The first strain, termed Degrader, degrades naphthalene into an intermediate metabolite, salicylate, while the second strain, termed Detoxifier, further metabolizes salicylate into pyruvate. (C) Schematic diagram of the Autonomist strain that is capable of degrading naphthalene completely. (D) Schematic diagram of the design of our evolution experiments. All the experiments started by equally inoculating the Degrader and Detoxifier strains to the minimum medium. In the condition of mutualism, naphthalene was supplied in every cycle, so the two strains in the consortium engaged in mutualistic cooperation throughout the evolution. Under competitive conditions, pyruvate was supplied in every cycle so both strains grow competitively. Under the fluctuating conditioind, naphthalene and pyruvate were alternately supplied in the passaging cycles, as a result of which the two strains alternately engaged in mutualism or competition growth in different cycles. This evolutionary process was conducted for ∼270 generations in each condition, to assess whether an autonomous genotype can evolve from the mutualistic consortium.

We performed our evolution experiment by consistently passaging the cultures of the consortium in three different environments of different carbon source supplements (Figure 1D): (1) naphthalene was supplied in every passaging cycle as the sole carbon source; as a consequence, the two strains within the consortium engaged in mutualistic cooperation throughout the evolution (referred to as mutualism condition thereafter); (2) pyruvate, which can be used by both strains for growth (Figure 1B), was supplied in every passaging cycle; as a consequence, both strains compete for the sole carbon source (referred to as the competition condition thereafter); (3) To mimic environmental fluctuations, naphthalene and pyruvate were alternately supplied in the passaging cycles; as a consequence, the two strains alternately engaged in mutualism or competition in different cycles (referred to as the fluctuating condition thereafter). we conducted 12 experimental replicates for each condition (i.e., 36 evolutionary lineages in total) to explore whether autonomous genotypes can evolve from the mutualistic consortium after the experimental evolution.

### Distinct evolutionary outcomes of the mutualistic consortium under different environmental conditions

To investigate the evolutionary fates of our mutualistic consortium under various conditions, we tracked the dynamics of the two strains in our evolutionary lineages and measured the phenotypic changes of the evolved populations, including their growth and tolerance to salicylate toxicity.

#### Degrader dominated the community when our mutualistic consortium evolved under the mutualism condition

In the mutualism condition, we observed that Degrader rapidly dominated all the lineages Figure 2A). After we ran the experiment for 280 generations in the mutualism condition, the Detoxifier strain was not detected, suggesting that the cooperation between Degrader and Detoxifier collapsed (Figure S3). When we cultured the evolved consortia using naphthalene as the sole carbon source, we found that these consortia exhibited significantly higher growth rates than ancestral mutualistic consortia (Student’s T-test, *p* < 0.05). However, their maximum cell densities (*OD*_*max*_) did not increase after evolution (Student’s T-test, *p* > 0.05; Figure 2B; Figure S4A). Next, we spread the cultures of these lineages on agar plates and picked four single colonies from each lineage. We confirmed that these isolated strains all evolved from the ancestral Degrader strains by sequencing their barcode tags (Table S1).

**Figure 2.**
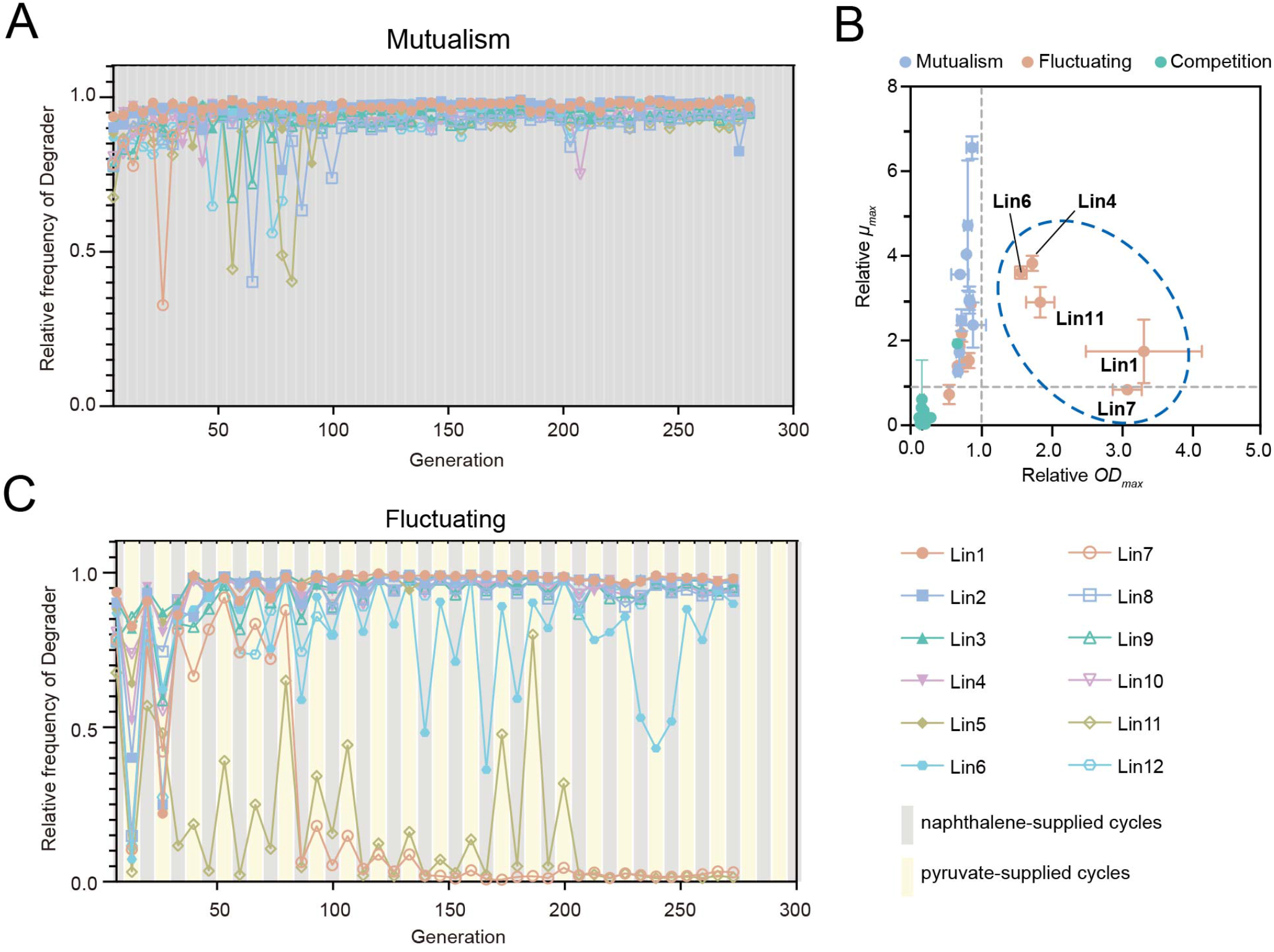
Dynamics of the mutualistic consortium. (A, C) The dynamics of the community composition (denoted by the relative frequency of the Degrader strain) across all the evolutionary lineages in the mutualism (A) and the fluctuating condition (C). 12 independent replicates (i.e., lineages) were performed for each condition, which are indicated by the curve legends shown on the right of (C). (B) Comparison of the growth kinetics between the evolved consortia and ancestral consortia. To determine their relative levels, we estimated maximum cell densities (*OD*_*max*_) and maximum growth rates (*μ*_*max*_) of the consortia evolved under each condition and those of the ancestral consortia (*n* = 3). Dashed lines indicate levels of the ancestral consortia. The five lineages that evolved under the fluctuating condition and exhibited higher growth kinetics to the ancestral consortia are circled and labeled.

When we cultured these isolated strains in minimum medium, we observed three distinct phenotypes in these isolated strains as compared to the ancestral Degrader strain. The first group of strains grew faster than the ancestral Degrader but their tolerance to salicylate toxicity did not change significantly (as surrounded by a blue oval in Figure S4B). The second group did not increase in growth rate but evolved to be more tolerant to salicylate toxicity (as surrounded by an orange oval in Figure S4B. The third group evolved to possess both higher growth rates and higher tolerance (as surrounded by a red oval in Figure S4B). Interestingly, we found that the first and second groups coexisted in lineage 6 (Figure S4B), suggesting that a novel form of niche differentiation between the two populations evolved after 280 generations in the mutualism condition. In addition, the third phenotype evolved and dominated in lineages 1 and 10 (Figure S4B), indicating that the ancestral Degrader strain was able to compensate for its deficiency (i.e., salicylate tolerance) by evolving itself. This observation parallels the findings from another study where the collapse of mutualistic cooperation was observed. In that study, a *Bacillus subtilis* strain deficient in biofilm formation was able to regain complete functionality and form a biofilm through self-evolution^54^. Similarly, in both our experiment and this previous study, one of the ancestral genotypes compensates for its lacking function by evolving a new function on its own, enabling it to thrive more effectively in its respective habitats. Together, these results demonstrated that the mutualistic cooperation between the Degrader and Detoxifier strains in the ancestral consortium collapses under the mutualism condition. Furthermore, we observed the evolution of either a novel form of niche differentiation or a single population with improved survival. However, we did not observe the evolution of an autonomous genotype that can perform the entire pathway of naphthalene degradation.

#### Divergent evolutionary outcomes of our mutualistic consortium in the competition condition

When passaging our DOL consortium under the competition condition, we observed that Degrader decreased in frequency during the early stages of the passage in all lineages, and this decrease persisted for approximately 100 generations in eleven of twelve lineages (Figure S5A). By performing competitive fitness assays, we found that this decrease was due to the higher fitness of Detoxifier in pyruvate-supplied environments than that of Degrader (Figure S5B-D). However, we found that the frequencies of Degrader of these lineages gradually exhibited significant variations after 100 generations, with the relative frequency of the Degrader strain ranging from 0.05 to 0.96 at the 268 generation. Specifically, the relative frequency of the Degrader strain exceeded the initial frequency of the inocula in six of the twelve lineages (Figure S5A). Together, these results suggested that the competition condition enabled the emergence of the Degrader mutants possessing varied fitness.

#### Autonomous genotypes evolved from the cooperative specialists under the fluctuating environment

We next investigated the evolutionary fates of our mutualistic consortium in the fluctuating condition. When we tracked the community dynamics, we found that the relative frequency of Degrader oscillated in most of the lineages (Figure 2C). Specifically, we found that while the Degrader strain exhibited higher frequencies in the naphthalene-supplied cycles, its frequency decreased when the cultures were passaged to the pyruvate-supplied cycles (Figure 2C). These results indicated that introducing the pyruvate-supplying cycles fosters the coexistence of both strains, in which they re-engage in competing for resources, ultimately preventing the collapse of their coexistence occurring in the mutualism condition. Interestingly, when we cultured the evolved consortia using naphthalene, we found that consortia from five of the twelve lineages exhibited higher maximum biomass yield than the ancestral mutualistic consortium, namely lineage No.1, No.4, No.6, No.7, and No.11 (Figure 2B; Figure S6A). We next spread the cultures of these five evolved consortia on agar plates and picked four single colonies from each lineage. We found all these isolates grew alone using naphthalene as their sole carbon source. Specifically, 14 out of 20 isolates showed similar or higher *OD*_*max*_ as the Autonomist strain and all isolates showed similar or higher growth rates as the Autonomist strain (Figure 3A, Student’s T-test, *p* < 0.05; Figure S6B).

**Figure 3.**
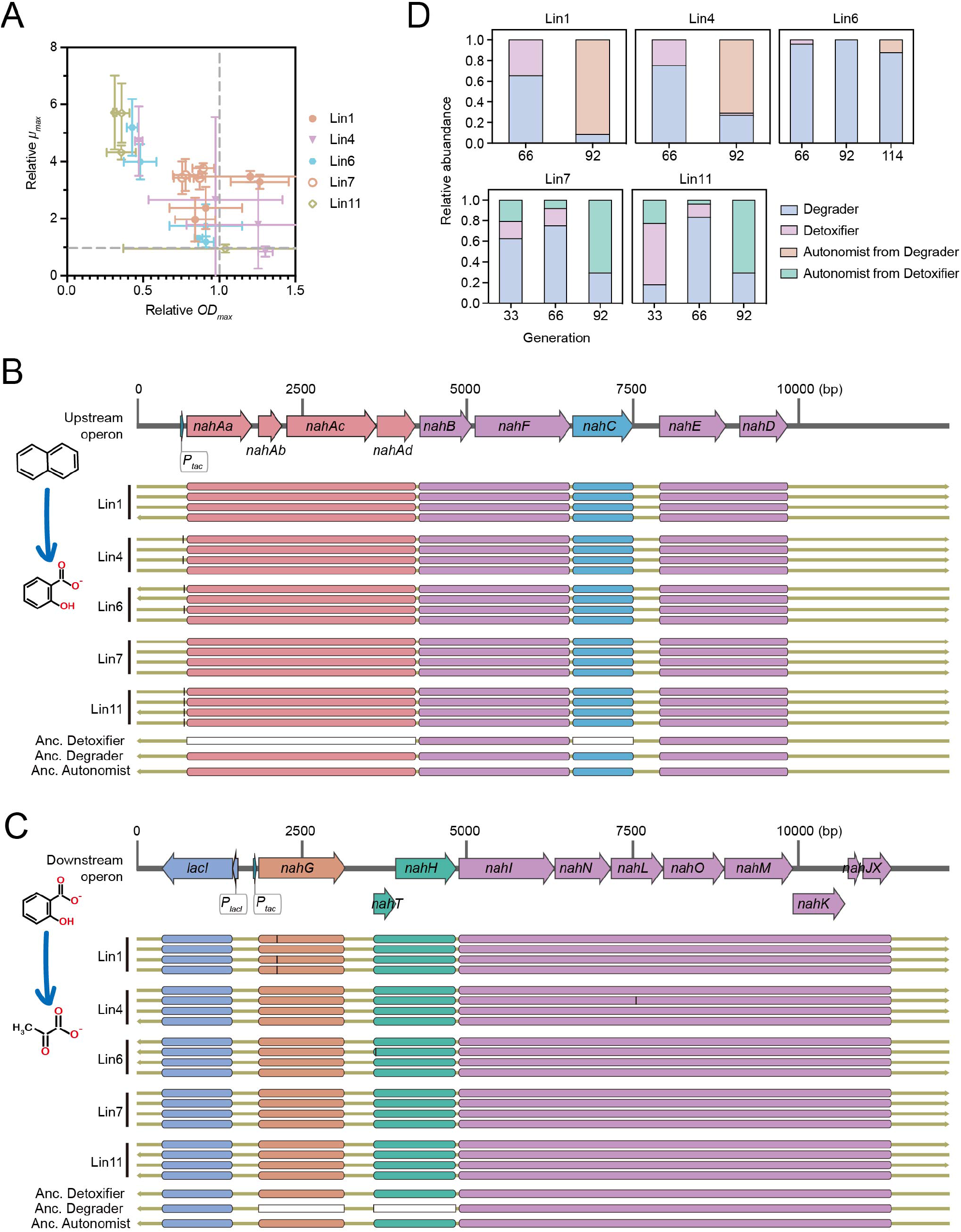
Autonomous populations evolved under fluctuating conditions. The five lineages (lineage 1, lineage 4, lineage 6, lineage 7, and lineage 11) that show approximate growth kinetics to the ancestral Autonomist (See Figure 2) are further analyzed. Four clonal isolates were picked from each evolved lineage. (A) Comparison of the growth kinetics between the evolved isolates and the ancestral Autonomist strain. To determine their relative levels, we estimated maximum cell densities (*OD*_*max*_) and maximum growth rates (*μ*_*max*_) of the evolved isolates and those of the ancestral Autonomist strain (*n* = 3). Dashed lines indicate levels of the ancestral Autonomist strain. (B-C) Comparative sequence analysis of two naphthalene operons in ancestral and evolved populations. The first operon (B) contains the genes responsible for the degradation of naphthalene to salicylate, while the second operon (C) encodes the further conversion of the salicylate to pyruvate. The ancestral Detoxifier strain lacks two key genes, *nahA* (Pink) and *nahC* (Blue) in the first operon, while the ancestral Degrader strain lacks *nahG* (Orange) and *nahTH* (Green) in the second operon. These mismatches are indicated by the white boxes. Through the whole-genome sequencing, we found all the evolved isolates regained these key genes. These regained genes are located exactly at the same genomic locus as that of our ancestral Autonomist strain. The gene sequences are exactly the same except for several single-base mismatches (denoted by the black line). The directions of the arrows indicate the direction of the sequenced contigs. (D) Tracing the emergence of Autonomist during evolution. Through the isolation of 24 single clones and PCR genotyping (See Methods), we measured the relative abundances of Degrader, Detoxifier, and the evolved Autonomist at two or three different generations in the five lineages in which Autonomist evolved under the fluctuating condition.

To explore the genetic underpinnings behind the improved growth of these isolated strains using naphthalene, we conducted whole-genome sequencing on all isolates. We first checked the genomic locations encoding the two operons of naphthalene degradation genes. Surprisingly, we found that all isolates evolved to regain a copy of the previously deleted genes required for naphthalene degradation (Figure 3B-C), potentially via HGT from its cooperative partner. Notably, these regained genes are located exactly at the same genomic locus as that of our ancestral Autonomist strain (Figure 3B-C), suggesting that the genes are regained via homogenous recombination^55,56^, one widespread mechanism benefiting HGT among different bacteria^57^. To eliminate the possibility that these observed Autonomist strains were derived from the contamination of the lab-stored Autonomist strain, we checked the sequence of the barcode tags. We found that the strains from three of the five lineages possessed the identical barcode tag as their ancestral Degrader strain, while the strains from two of the five lineages possessed the identical barcode as their ancestral Detoxifier strain (Table S2). This result clearly suggests that the observed Autonomist strains evolved from either Degrader or Detoxifier during the process of strain evolution. Together, these results indicated that the ancestral cooperative specialists in our mutualistic consortium evolved into autonomous genotypes, potentially driven by HGT. Nevertheless, this evolution only occurred in the fluctuating condition where naphthalene and pyruvate were alternately supplied.

### Fluctuating environments create conditions critical for the evolution of autonomy

To explain why the fluctuating environment is mandatory for the evolution of autonomy in our experiment (in other words, why both mutualism and competition cycles are required), we proposed three hypotheses and performed corresponding analyses to assess their validity.

First, we hypothesized that the mutualism cycles are crucial to exert selective pressure in favor of Autonomist over mutualistic cooperation. In our experiments, we observed that the evolved Autonomist strains exhibited faster growth and higher maximum biomass yield compared to the mutualistic consortium when grown on naphthalene as the carbon source (Figure 1A). However, this growth advantage was not observed in the medium supplemented with pyruvate (Figure S5). Furthermore, we observed that the evolved genotypes under competition conditions, where pyruvate served as the sole carbon source in all cycles, exhibited a lower capability for naphthalene degradation (Figure S7A-B) and the autonomous genotype did not evolve in this condition (Figure S7C). This result indicated that the competition cycles alone did not favor the evolution of the autonomous genotype. Together, these findings support our hypothesis and suggest that the mutualism cycles are therefore necessary for the evolution of autonomy in our experiments.

Second, we hypothesized that competition cycles enabled the better coexistence of Degrader and Detoxifier, allowing the evolution of Autonomist via HGT. In our experiments, we observed that Degrader and Detoxifier exhibited a low level of coexistence in cycles supplemented with naphthalene, with Degrader dominating. However, in cycles supplemented with pyruvate, we observed a significant improvement in their coexistence (Figure 2C). As shown in Figure S6, this increase in coexistence is attributed to the higher fitness of Detoxifier in pyruvate-supplied cycles than Degrader, resulting in an increase in its frequency and facilitating its coexistence with Degrader. Although we observed that Degrader may evolve to possess higher fitness under the competition conditions, our results suggested that Autonomist evolved at an earlier stage, spanning 33 to 114 generations that precede the stage where Degrader evolved to higher fitness (Figure 3D). The increased coexistence is expected to benefit the evolution of Autonomist via HGT because the coexistence of the donor and recipient is the prerequisite for HGT^30^. To further examine the importance of co-existence, we performed evolution experiments on twelve novel lineages under the mutualism condition.

Although Degrader and Detoxifier showed low co-existence levels under this mutualism condition, we manually re-added new Detoxifier cells to the cultures to increase the co-existence. We found that Autonomist evolved in three of the twelve lineages (Figure 4). This result confirmed that the better coexistence of Degrader and Detoxifier in the competition cycles plays a key role in driving the evolution of Autonomist via HGT.

**Figure 4.**
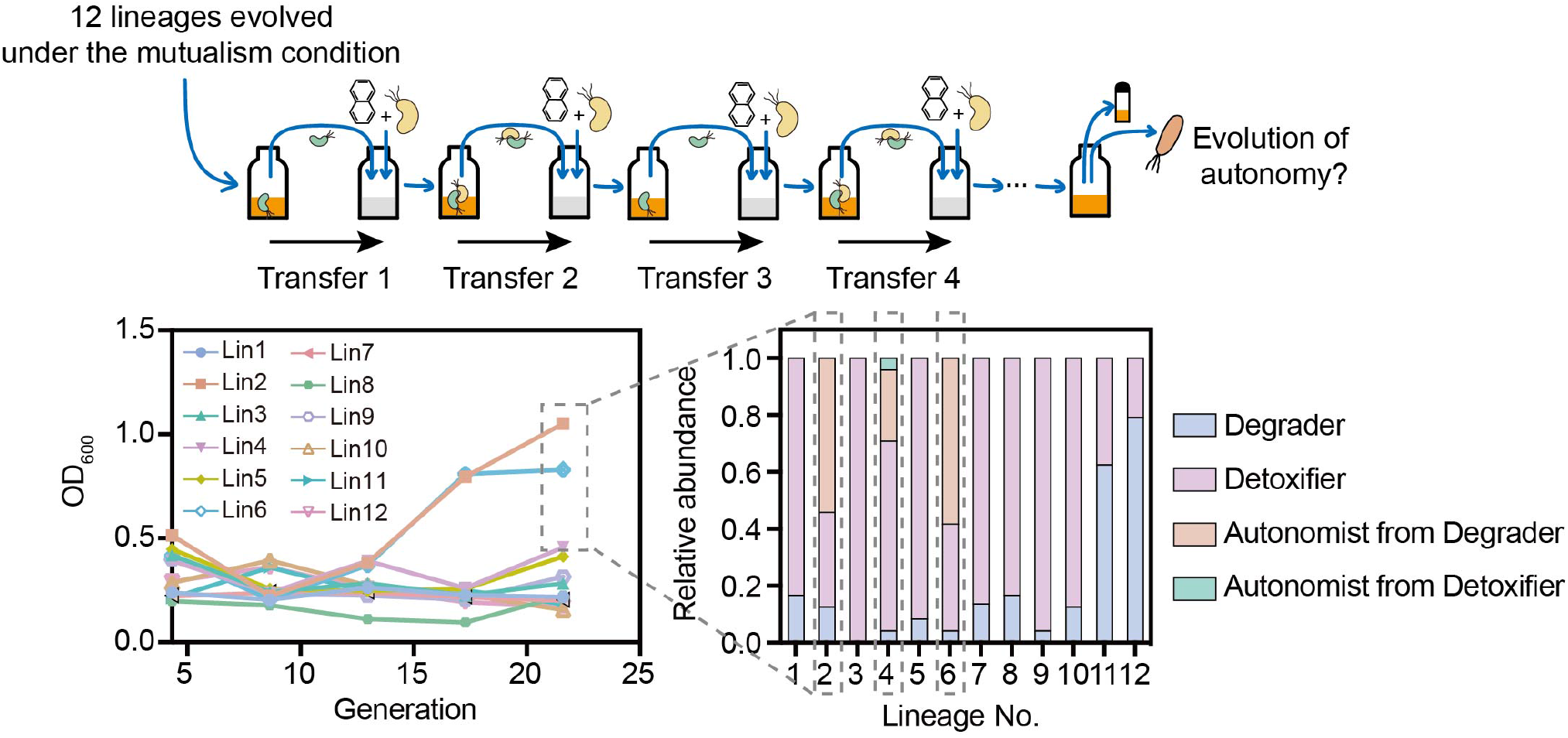
Supplementing the Detoxifier strain to Degrader-dominated consortia favors the evolution of the Autonomist strain. After our initial evolution experiments, the Autonomist strain did not evolve in all twelve lineages under the mutualism condition. We then started a new evolution experiment of the ancestral consortia by reintroducing the Detoxifier strain when transferring the cultures to fresh media. The Detoxifier strain was reintroduced to the twelve lineages pre-evolved under the mutualism condition. After 22 generations, reintroducing the Detoxifier strain enabled the evolution of the Autonomist strain in three out of twelve lineages.

Third, the mutualistic consortium accumulates higher cell density in the pyruvate-supplied cycles (OD_600_: ∼1.0) than that in the naphthalene-supplied cycles (OD_600_: ∼0.4; Figure S8). Previous studies reported that higher cell density increases HGT rates by increasing the chance of cell-cell contact^58^, or the secretion and acquisition of extracellular DNA^59,60^. Thus, the increase in cell density in the pyruvate-supplied cycles may increase the HGT probabilities between Degrader and Detoxifier, and further benefit the evolution of Autonomist.

### Mathematical modeling of the evolution of autonomy

#### Quantitative explanations for the evolution of autonomy from the mutualistic consortium under fluctuating conditions

To gain quantitative insights into why the fluctuating condition led to the evolution of autonomy in our experiments, we developed a mathematical model grounded on three key assumptions derived from our experimental observations. First, we considered two growth environments: (1) An environment that allows for the facultatively mutualistic cooperation between Degrader and Detoxifier (referred to as environment (Mutual.) thereafter), where Degrader degrades a substrate into a toxic intermediate, and Detoxifier further metabolizes this intermediate for detoxification (Figure S9A). (2) An environment where both strains compete for limited resources (referred to as environment (Comp.) thereafter). Second, Autonomist can evolve from any of Degrader and Detoxifier via HGT. We assumed that the HGT probability positively correlates with the cell density and the co-existence level of Degrader and Detoxifier^61,62^ (See Methods). Third, we assumed that the evolved autonomous genotype possesses a growth advantage (in terms of both growth rate and biomass yield) in the environment (Mutual.); not, however, in the environment (Comp.). Based on these assumptions, we built an ODE system (Methods: Eqn. [2] to [10]) to mimic two important conditions in our experiments: (1) the mutualism condition where the initial consortium is passaged continually in the environment (Mutual.); (2) the fluctuating condition in which the consortium grows alternately in the environment (Mutual.) and the environment (Comp.).

To reproduce the basic findings of our experiments, we performed simulations initialized with the parameters obtained from our experimental system (Table S3: default values). Out of 100 replicated simulations of the fluctuating condition, we found that Autonomist evolved in 59 replicates within 50 dilution cycles (Figure 5A-C); in comparison, Autonomist only evolved in 16 of 100 replicates in the mutualism condition (Figure 5D-E). This result agrees with our experimental findings, suggesting that environmental fluctuations facilitate the evolution of autonomy within the conditions of our experiments.

**Figure 5.**
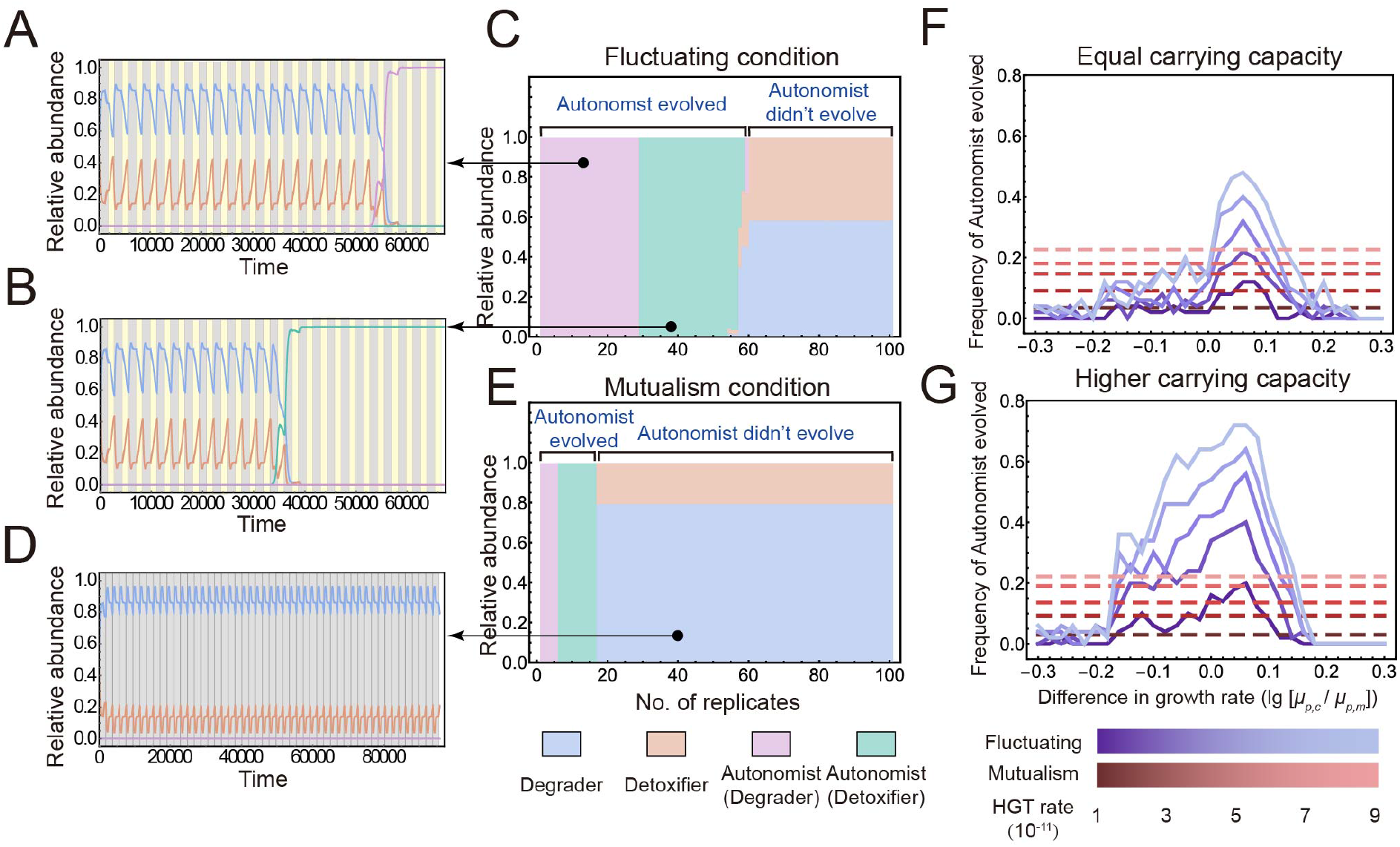
Representative simulations using our mathematical model capture the basic findings of our experiments. The simulations were initialized with the parameters measured from our experiment systems (Table S3: default values). Simulations under fluctuating and mutualism conditions were performed. (A-B) the relative abundance of different genotypes over the two representative simulations under the fluctuating condition, in which Autonomist evolved from Degrader (A) or Detoxifier (B). (C) The summary of the evolutionary outcomes of 100 independent replicates was conducted for the fluctuating condition. (D) the relative abundance of different genotypes over a representative simulation under the mutualism condition. (E) The summary of the evolutionary outcomes of 100 independent replicates was conducted for the mutualism condition. (F-G) The correlation between the relative growth rate of the Detoxifier strain to that of the Degrader strain in the environment (comp.) and the frequency of the evolution of autonomy was analyzed under two scenarios: (F) when the maximum carrying capacities of the two environments were the same; (G) the carrying capacity of the environment (Comp.) was set to be twice that of the environment (Mutual.). Unless specified otherwise, all these simulations were parameterized by the default values shown in Table S3.

Our experiments suggested two important hypotheses regarding how the environment (Comp.) in the fluctuating conditions select for the evolution of autonomy via HGT: (1) the higher fitness of Detoxifier in the environment (Comp.) increases the co-existence of both strains and thus benefits the HGT; (2) the higher cell density in the environment (Comp.) facilitates the evolution of autonomy. To quantitatively examine the first hypothesis, we set the carrying capacity of the two environments to be identical (to eliminate the effects of any differences in cell density). We performed simulations parameterized with varied relative growth rates between Degrader and Detoxifier. The simulation results showed that the evolution of autonomy was favored under the fluctuating condition when the growth rate of Detoxifier fell within a specific range: higher than that of Degrader but lower than a threshold value (Figure 5F). Our analysis showed that this range closely corresponded to the range in which the coexistence level of the two strains was higher under the fluctuating condition compared to the mutualism condition (Figure S9B). These results provide compelling evidence that when the fitness of Detoxifier within the competitive environment (Comp.) falls within our defined range, the coexistence of the two strains increases, which further promotes the probability of HGT. These findings robustly support our first hypothesis.

To test our second hypothesis as to whether the higher cell density in the environment (Comp.) facilitates the evolution of autonomy, we performed additional simulations. We set the carrying capacity of the environment (Comp.) to be twice that of the environment (Mutual.) to enable higher cell densities in the environment (Comp.), in line with the experimental setting of our evolution experiment. Under this setting, we observed an expansion in the value range of Detoxifier fitness that leads to an increased probability of HGT and the subsequent evolution of autonomy (Figure 5F-G). Furthermore, we found that this range can be predicted by combining the effects of cell density and the coexistence level of the two strains (Figure S9B-C). Together, the findings of these simulations demonstrated that environmental fluctuations play a crucial role in introducing variations in relative fitness and cell density, thereby facilitating the evolution of autonomy through HGT within specific conditions.

#### The effect of fluctuating environments on the evolution of autonomy is pathway-dependent

To test the general validity of our experimental findings, we performed simulations using 17501 randomly-sampled parameter sets, each of which defined a specific condition of pathway kinetics and strain growth (Methods; Table S3). For a given set, we modeled both the mutualism condition and the fluctuating condition with 20 replicates for each, which allowed us to calculate the frequency of the evolution of autonomy in both conditions (defined as *P*_*mut*_ and *P*_*fluc*_). As shown in Figure 6A, we observed that Autonomist evolved in at least one of the two conditions in the simulations conducted with 13905 parameter sets, accounting for nearly 80% of all simulated sets. Notably, 2269 of these sets (∼13%) resulted in a scenario closely resembling our experimental observations, in which the autonomous genotype exclusively evolved in the fluctuating condition. In comparison, a smaller number of sets (1003, ∼6%) enabled the simulations where the autonomous genotype only evolved in the mutualism condition. Moreover, among the simulations where the autonomous genotype evolved in both conditions (10633 of 17501 sets, ∼61%), we observed a significantly higher frequency of the mutualism consortium evolving into an autonomous genotype in the fluctuating condition compared to the mutualism condition (Figure 6B; *p* < 0.001). Together, these results indicated that the influence of fluctuating environments on the evolution of autonomy is dependent on the specific conditions of the pathways involved.

**Figure 6.**
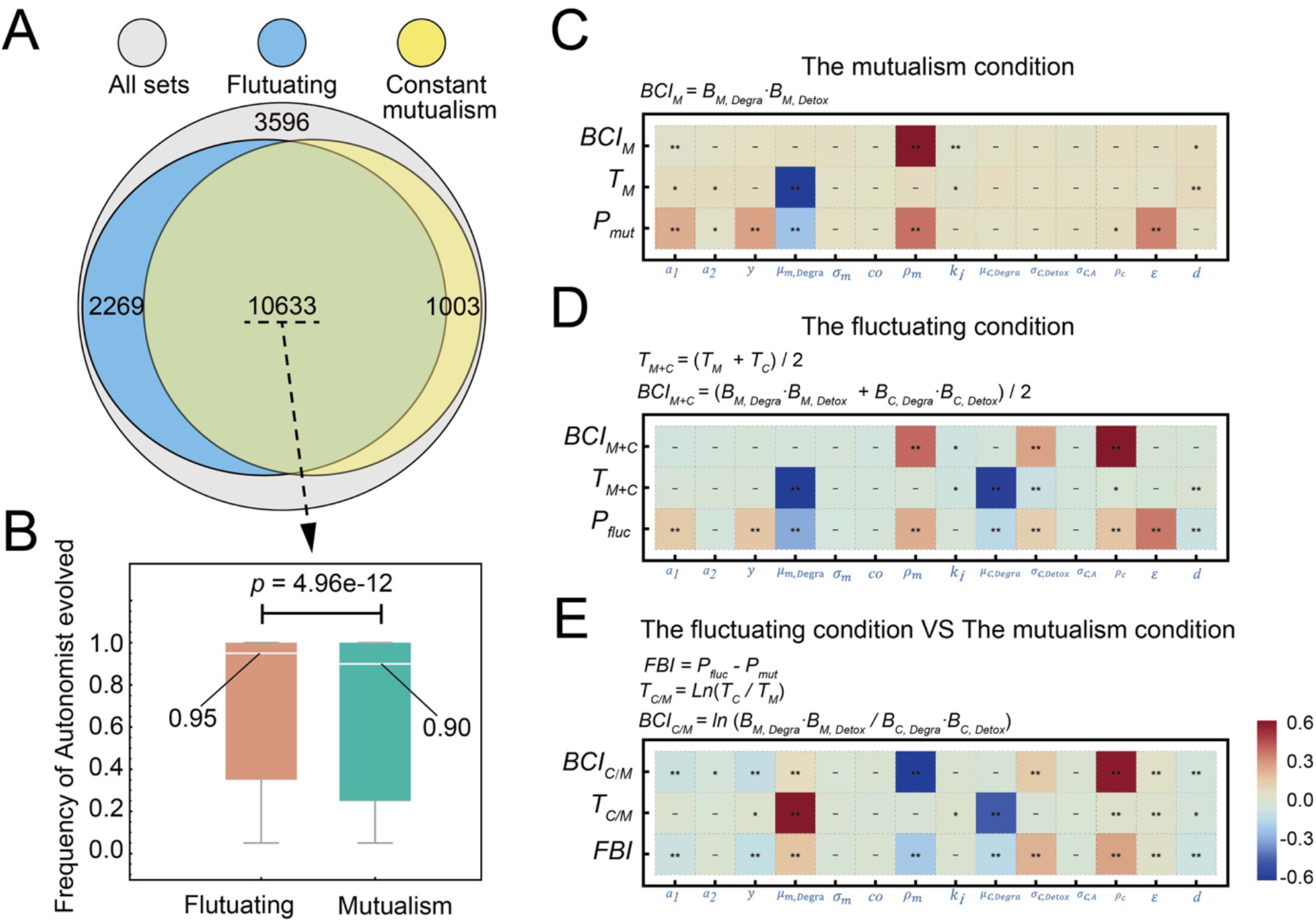
Mathematical model of the evolution of autonomy from cooperative specialists. We performed simulations using 17501 parameter sets that are generated based on random sampling from given value ranges (Table S3). For a given set, we modeled both the mutualism condition and the fluctuating condition with 20 replicates for each and calculated the frequency of Autonomist evolution in both conditions (defined as *P*_*mut*_ and *P*_*fluc*_). (A)Venn diagram summarizing the evolutionary outcomes of all simulations, indicating whether Autonomist evolved in both conditions (green), in either one of the conditions (blue and yellow), or in neither condition (Gray). (B) Comparison of the *P*_*mut*_ and *P*_*fluc*_ of the simulations where the autonomous genotype evolved in both fluctuating and mutualism conditions (the green section in (A)). The median values were indicated and pairwise Student’s T-tests were performed to examine their significant difference. (C-E) Key factors determining the probability of the evolution of autonomy under the mutualism condition (C) and the fluctuating condition (D), and key factors determining whether the fluctuating condition benefits the evolution of autonomy (E). We performed Pearson correlation analysis between the parameter values and the frequency of Autonomist evolution (*P*_*mut*_ *P*_*fluc*_ and *FBI*), cell density and coexistence level in the two environments (*BCI*_*M*_, *BCI*_*C*_, and *BCI*_*C/M*_), as well as duration of growth in the two environments (*T*_*M*_, *T*_*C*_, and *T*_*C/M*_). The Pearson correlation coefficients are indicated by the color density; the statistical significance of the analysis is indicated as * *p* < 0.01; ** *p* < 0.0001. Our results also show a positive correlation between *P*_*mut*_ and the *BCI*_*M*_ value (R^2^ = 0.28; *p* < 0.001) under the mutualism condition, and a positive correlation with the average values of both *BCI*_*M*_ and *BCI*_*C*_ under the fluctuating condition (R^2^ = 0.39; *p* < 0.001), suggesting that higher cell density and coexistence level benefit the evolution of Autonomist.

#### Key factors determining the evolution of autonomy from mutualistic cooperation

To investigate the key factors defining whether Autonomist can evolve in the mutualism condition or the fluctuating condition, we compared the distributions of parameter values that enabled the evolution of autonomy in simulations to those factors that lacked this property. We identified five of 14 parameters that significantly affected whether Autonomist can evolve under the mutualism condition (Figure S10; Kolmogorov-Smirnov Test: *p* < 0.01). This finding was further supported by our Pearson correlation analysis between the parameter set and the frequency of the evolution of autonomy in the corresponding simulations (Figure 6C; correlation coefficients over 0.1). Using identical methods, we identified eight of 13 parameters that significantly affect whether Autonomist can evolve under the fluctuating condition (Figure S11; Kolmogorov-Smirnov Test: *p* < 0.01; Figure 6D). Furthermore, our analysis suggested that *ρ*_*m*_ (maximum carrying capacity in the environment (Mutual.)) positively affected cell density and coexistence levels under the mutualism condition (quantified by *BCI*_*m*_, see Equation [11]), thereby facilitating the evolution of autonomy (Figure 6C). Similarly, *ρ*_*m*_, *ρ*_*c*_ (maximum capacity in the environment (Comp.)), and *σ*_*c,Detox*,_ (growth rate of Detoxifier relative to that of Degrader in the environment (Comp.)) positively impacted cell density and coexistence levels under the fluctuating condition (quantified by *BCI*_*M+C*_), consequently benefiting the evolution of autonomy (Figure 6D). We also found that *μ*_*m*_, _*Degra*_ (Growth rate of Degrader in the environment (Mutual.)) and *μ*_*c,Degra*_ (Growth rate of Degrader the environment (Comp.)), respectively affected the evolution by modifying the duration of each cycle (Figure 6C-D). Higher growth rates shorten the cycle duration (because we passaged the consortium once they reached the stationary phase), thus reducing the probability of HGT. Lastly, we analyzed the parameters defining when the fluctuating condition benefits the evolution of autonomy compared to the constant mutualism condition. We hypothesized that environmental fluctuations are beneficial, providing that the mutualistic consortium exhibits a higher cell density and better coexistence of the two strains in the environment (Comp.) compared to the environment (Mutual.). To test this hypothesis, we conducted a correlation analysis between a Relative Biomass-Coexistence Index (*BCI*_*C/M*_; Methods) and a Fluctuation-Benefiting Index (*FBI* ; evaluating the degree to which fluctuating conditions benefit the evolution of autonomy; Methods). Our analysis revealed a significant positive relationship between *BCI*_*C/M*_ and *FBI*, supporting our hypothesis (*p* < 0.001; R^2^ = 0.41). We also found a positive correlation between the relative duration of growing in the environment (Comp.) versus the environment (Mutual.) (Refer to as *T*_*C/M*_ thereafter) and FBI (*p* < 0.001; R^2^ = 0.35), suggesting that longer duration in the environment (Comp.) also benefit the evolution of autonomy. Next, we used Pearson correlation analysis to identify key parameters that largely affect *BCI*_*C/M*_ and *T*_*C/M*_ (Figure 6E): (1) *ρ*_*m*_ negatively influences the values of *BCI*_*C/M*_ while *ρ*_*c*_ and *σ*_*c,Detox*,_ (growth rate of Detoxifier relative to that of Degrader in the environment (Comp.)) affects *BCI*_*C/M*_ positively. (2) *T*_*C/M*_ is positively impacted by *μ*_*m,Degra*_ while negatively altered by *μ*_*c,Detox*_. In summary, our findings suggest that the evolution of autonomy in the mutualistic consortium is promoted by a fluctuating condition when the consortium exhibits increased cell density, enhanced co-existence, and prolonged growth in the environment (Comp.). These factors are determined by the relative growth rate and capacity in the two environments.

## Discussion

Here, we provided experimental evidence that a consortium engaged in mutualistic cooperation can evolve into an autonomous genotype via horizontal gene transfer (HGT). We show experimentally that environmental fluctuations are important to drive such evolution and mathematically identify the key parameters that determine when fluctuating environments are beneficial.

In his famous work ‘Wealth of Nations’, Adam Smith proposed that cooperation emerges from autonomy thus achieving higher productivity^63^. This concept originally proposed for economic mechanisms has later been applied to better understand the evolution of biological systems. Many studies have since discussed when and how cooperation evolves from an autonomous ancestor^2,64-68^. However, autonomous organisms are favored in many specific environments over cooperation in nature. Therefore, it is also crucial to understand when and how autonomous organisms can evolve from cooperation^16,69,70^. Such evolution is particularly critical in scenarios where cooperation initially evolved in an environment that favored cooperation, but later shifted to favor the autonomous organisms. Several studies predicted that genetic exchange (e.g., HGT) among natural partners may help a cooperative specialist regain deficient function, which could mediate the collapse of cooperation and the evolution of autonomy. Here in this study, we provide experimental evidence that supports this hypothesis. A recent review suggested that mutualisms exhibiting a lower degree of dependency, such as facultative mutualisms, are more likely to transition back to autonomy compared to those that show higher dependency, such as obligate mutualisms^71^. Given that the two cooperative specialists in our consortium display facultative mutualism, our results provide experimental evidence supporting this prediction.

Our study demonstrates that environmental fluctuations favor such evolution. Since natural populations often experience fluctuations in nutrient types in their environment^36^, we expected that this effect would be prevalent in nature. The underlying mechanism of our findings is that environmental fluctuations impact both the selection pressure on the autonomous genotype and conditions for HGT (i.e., high cell density and the coexistence of both genotypes). Based on our analysis, we emphasize how important coexistence is for driving these co-evolutionary pathways that result in the emergence of novel genotypes. In our case, to enable the sharing of their genetic elements via HGT, different genotypes must co-exist both spatially and temporally^30,35,59^. In another study, two mutualistic *E*.*coli* partners evolved to form multicellular spatial clusters that benefit their cooperation^72^. Similarly, a recent study showed that *Pseudomonas aeruginosa* evolved to increase resistance towards beta-lactam antibiotics when coexisting with *Staphylococcus aureus*^73^. In line with our case, the evolution observed in these studies required the coexistence of both partners. Therefore, co-existence allows genetic exchange and group-level selection that drives novel evolutionary events.

The idea to increase the co-existence of different partners also has implications for the artificial selection of high-function microbiomes^74-76^, in which the co-existence of different organisms must be maintained to enable the evolution to achieve the higher function of the community toward specific industrial uses. Our work offers a potential approach that manipulating the fluctuations of nutrient supply can increase species co-existence. However, fluctuating environments are not always useful for maintaining co-existence. For example, one recent study reported that as a result of the rapid evolution of one partner in a fluctuating environment, the long-term co-existence of two species engaged in positive pairwise interactions collapsed^77^. Our mathematical model can be used to quantitatively predict whether this method is efficient in a specific microbial system. Another method to select high-function microbiomes suggested by our experiments is to re-add the seeding species, which mimics the immigration events in nature^78^. As re-adding Detoxifier cells into our consortium also enables the evolution of autonomy, re-supplying the key seeding species to the other microbial systems may increase their function.

We also observed randomness in our experiments: the evolution of autonomy in only five of the twelve lineages. We speculate that several stochastic factors are responsible for this outcome. First, the occurrence of HGT events is random, which requires occasional cell-cell contacts or the acquisition of extracellular DNA^60,79^. Second, the co-existence of both genotypes and the expansion of an autonomous genotype are also affected by random factors, including the bottleneck effects occurring during the culture dilution at the beginning of each cycle^80,81^, as well as phenotypic variations of the seeding cells among different lineages. In summary, the successful evolution of autonomy requires that the HGT events occur before the co-existence of two genotypes collapses and that the evolved autonomous genotypes can overcome other random factors to be successfully fixed in the community. Similar phenomena were observed in other studies^82^. For instance, while *E. coli* usually outcompetes *Saccharomyces cerevisiae* in co-culture, one study shows that a small number of lineages of *S. cerevisiae* evolved and coexisted, namely by mutating multi-hit genes before *S. cerevisiae* were excluded by *E. coli*^83^. Therefore, the chronological order of evolutionary events and the loss of coexistence, determined by stochastic factors, is key to shaping the outcomes of co-evolution. To capture the effects of such stochastic factors, further evolution experiments of microbial communities should include more replicates of experimental lineages.

In conclusion, our study addressed the knowledge gap regarding whether autonomy can evolve from mutualistic cooperation through genetic transfer. Our findings advance our understanding of evolutionary adaptations in biological systems, demonstrating their capacity to adapt to complex environments by flexibly switching their survival strategies. Furthermore, we offer fresh insights into directing the co-evolution of different species towards high-function artificial systems through manipulating environmental fluctuations.

## Methods

### Construction of the strains used in this study

The strains and plasmids used in this study are summarized in Table S4. We used a previously developed experimental microbial system for naphthalene degradation, which is described elsewhere^42^. Briefly, the Autonomist strain can degrade naphthalene to the final products (pyruvate and acetyl-coenzyme A). The Degrader strain contains a deletion in *nahG* (encoding a salicylate hydroxylase) and *nahTH* (encoding a catechol 2, 3-dioxygenase) genes. As a result, Degrader only oxidizes naphthalene to salicylate, during which byproduct pyruvate is generated thus supporting its growth. The Detoxifier strain contains a deletion in *nahA* (encoding a naphthalene dioxygenase) and *nahC* (encoding a 1, 2-dihydroxynaphthalene dioxygenase) genes, which only convert salicylate to the final products.

To distinguish ancestral genotypes during evolutionary experiments, ancestral Autonomist, Degrader, and Detoxifier were engineered that carry unique DNA barcodes (Table S1-2). To measure their relative abundance in the consortia, Degrader and Detoxifier were labeled with *mCherry* and *eGFP*, encoding red and green fluorescent proteins. To achieve the fluorescent labeling, the DNA fragments of the *barcode* and *mCherry* gene, or the *barcode* and *eGFP* gene, were cloned into a vector, pUC18T-mini-Tn7T-Gm (Table S4). The derived vector was delivered to host cells via conjugative four-parental mating^84^. As a result, these fragments were knocked into the chromosomes of these strains at Tn7 attachment (attTn7) sites^84^. *P. stutzeri* exconjugants were selected in solid LB supplemented with nalidixic acid (5 *μ*g/mL) and gentamycin (25 *μ*g/mL). To prevent contamination during the evolution experiment, FRT excision of the gentamycin resistance marker was not performed, and all of the ancestral strains retained gentamycin resistance.

To observe the evolution of consortia over a relatively short time, the mutation rates of strains were increased by regulating the expression of the *mutS* gene (Figure S12). The promoter *rhaSR-PrhaBAD* induced by L-rhamnose was inserted in front of the *mutS* gene. During the evolutionary experiments, L-rhamnose was not added to the medium, and *mutS* was not expressed, leading to a higher spontaneous mutation rate due to the inactivation of the methyl-directed mismatch repair system^85^. After the evolutionary experiments, bacteria grew in the presence of L-rhamnose (0.2% w/v) for DNA extraction and preparation of growth curve measurement experiments. The expression of *mutS* resulted in a lower spontaneous mutation rate. This genetic modification was implemented by allele exchange using the suicide plasmid pK18mobsacB^86,87^.

### Salicylate tolerance assays

To measure the tolerance ability of ancestral and evolved strains to salicylate, overnight cultures were diluted and incubated with an initial OD_600_ (optical density at 600 nm) of 0.05 in fresh minimal medium supplemented with 5 g/L pyruvate and 40 C-mM salicylate along with no-salicylate negative control. Bacterial cultures were grown at 30°C with shaking at 200 rpm. OD_600_ was measured when the bacteria entered the stationary phase. The tolerance ability of bacteria to salicylate was determined as the percentage increase or decrease compared to when bacteria were grown in the absence of salicylate.

### Cultural Condition and the evolution experiments

Bacterial cultures were grown at 30°C on an orbital shaker set to 200 rpm. Pre-cultures were prepared by cultivating a single isolated colony from a lysogeny broth (LB) plate in the rich medium^88^ (Yeast extract 10 g/L, beef extract 5 g/L, peptone 10 g/L, ammonium sulfate 5 g/L) until late-exponential phase (∼12 h). Pre-cultured cells were centrifuged and washed three times with the minimal medium^89^ prior to the start of the experiments. The seed of the synthetic consortium was made by equally mixing the Degrader strain and the Detoxifier strain. Then the cultures were inoculated with an initial OD_600_ of 0.05 and performed in the minimal medium supplemented with pyruvate (0.5% w/v) or naphthalene powder (1% w/v) as the sole carbon source. The cultures were performed in 20-mL scintillation containing 4 mL of the medium. The evolution experiments were performed under three different conditions (Figure 1). (1) the mutualism condition. The consortium was propagated through 65 growth-dilution cycles in the minimal medium supplemented with naphthalene as the sole carbon source. The consortium was initially transferred every three days for a total of 24 transfers, followed by 41 transfers every two days. The cultures were diluted by a factor of 20 into fresh media during each transfer (∼4.3 cellular divisions per growth cycle). The experiments lasted for a total of 154 days, corresponding to approximately 280 generations of bacterial growth. (2) the competition condition. The consortium was propagated through 40 growth-dilution cycles by daily (24 h) transfers in the minimal medium supplemented with pyruvate. During propagation, cells were diluted by a factor of 100 into fresh media, which corresponded to ∼6.7 cellular divisions per growth cycle and ∼268 cellular divisions over a growth cycle. (3) The fluctuating condition. The culture was propagated by switching between the minimal medium supplemented with naphthalene and the medium supplemented with pyruvate. The naphthalene-supplied cycles lasted for three days at the first 17 cycles, following two days for the rest eight cycles. The pyruvate-supplemented cycles lasted for one day. The cultures were diluted by a factor of 100 into fresh media during each transfer (∼6.7 cellular divisions per growth cycle). In total, 41 transfers were performed (∼275 generations).

In total, 36 replicated lineages were experimentally evolved, in which 12 independent lineages were evolved in parallel for each condition. Biomass and relative abundances of the two populations were assessed by measuring OD_600_ and fluorescence intensity via spectrophotometry in a plate reader. During evolution, glycerol stocks (20% glycerol) were prepared every five or six transfers and stored at -80°C for future analysis. In the propagation of the mutualism condition, two out of twelve lineages were excluded from further analysis as the two lineages went extinct after 50 generations.

At the end of the propagation experiment, the evolved consortia were spread on LB agar plates supplemented with 0.2% (w/v) L-rhamnose to isolate evolved clones. To determine whether Autonomist evolved in the cultures and to measure the relative abundances of different genotypes, the four key genes (*nahA, nahC, nahG*, and *nahTH*) encoding naphthalene degradation, a subset of which were deleted in Degrader and Detoxifier, were amplified by PCR to detect their presence. If all four genes were detected, the clones were identified as the putative evolved Autonomist; If only *nahA* and *nahC* were detected, the clones were identified as Degrader; If only *nahG* and *nahTH* were detected, the clones were identified as Detoxifier. For each lineage, 24 clones were randomly picked and tested by PCR for each lineage. Next, four putative Autonomist colonies from each lineage were selected for growth kinetics measurement and whole-genome sequencing. The isolates were individually grown overnight in MF medium supplemented with 0.5% (w/v) sodium pyruvate and 0.2% (w/v) L-rhamnose and stored in glycerol (final concentration 20%) at -80°C until further use.

### The evolution experiments of re-introducing Detoxifier and examination of the evolved Autonomist

An experiment was designed to further test whether the coexistence of Degrader and Detoxifier is the key determinant for the evolution of Autonomist. After our initial evolution experiments, Autonomist did not evolve in all twelve lineages under the mutualism condition. A new experimental evolution of the ancestral consortium was performed, in which Detoxifier was re-added to the culture before each passage (final OD_600_ = 0.025, approximately 2.5×10^7^ cells). The cultures were passaged for five cycles under the mutualism condition. The same methods as shown in the above section were applied to determine whether Autonomist evolved in the cultures and to measure the relative abundances of different genotypes after evolution.

### Flow cytometry

Flow cytometry was performed with a CytoFLEX S (Beckman, USA) flow cytometer. During the coculture of the ancestral Degrader and Detoxifier, genotypes in the cultures were distinguished according to their respective fluorescent labeling (eGFP or mCherry). eGFP fluorescence was detected using a 488 nm excitation laser combined with a FITC (525/40 nm filter) detector. mCherry fluorescence was measured using a 561 nm laser combined with an ECD (610/20 nm filter) detector. Before the measurements, each sample was diluted to 10^−4^ - 10^−6^ cells/mL in saline solution. Samples were processed at a fixed flow rate of 30 µL/min for an acquisition time of 1 min. Flow cytometry results were analyzed using CytExpert software.

### Growth kinetics of ancestral and evolved consortia and clones

Aliquots (0.1 mL) of ancestral and evolved glycerol stocks were revived in 4 mL of the minimal medium containing 0.5% sodium pyruvate and 0.2% L-rhamnose and incubated at 30°C with shaking at 200 rpm. These overnight cultures were washed three times with MF medium. Then, cells were inoculated with an initial OD_600_ of 0.05 and were grown at 30°C with shaking at 200 rpm. Growth curves were measured by analyzing changes in cell density (OD_600_) at regular time intervals, and fitted by the following logistic function^90^:

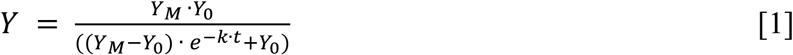

where *Y*_9_ is the minimum population biomass; *Y*_*M*_ is the maximum population biomass; and *k* quantifies the curve’s steepness. The following demographic parameters were extracted: the maximum predicted value for OD_600_ (Growth yield; unit, OD_600_) and the maximum rate of biomass increase (Growth rate; unit, h^-1^).

### PCR genotyping

To identify the genotypes of evolved consortia and isolates, the evolved consortia and isolates were subjected to PCR amplification using a mix of primers including four primer pairs (Table S5). Every PCR primer pair was designed to only amplify a product when the genome of the sample carried the specific gene sequence. The amount of PCR products depends on the kinds of genes in the sample. In addition, the four amplification products displayed different sizes, which can be distinguished into different bands by DNA electrophoresis. Degrader possessed two genes distinct from Detoxifier (*nahA* and *nahC*). Detoxifier possessed two genes (*nahG* and *nahTH*) distinct from Degrader. Autonomist possesses these four genes. The genotypes of the sample were identified by detecting the presence or absence of the four genes.

### Whole-genome sequencing and data analysis

In total, 136 clones were picked from the evolved populations (four clones per lineage; 12 lineages for the competition condition and the fluctuating condition, and 10 lineages for the mutualism condition). These clones were grown to saturation in RB media supplemented with 0.2% L-rhamnose and 25 *μ*g/mL gentamicin. 1 mL of overnight culture was used to extract DNA. Sequencing library preparation was done by using the DNA library preparation kit (Illumina). DNA sequencing was performed on Illumina Novaseq 6000 with 150 bp paired-end. As the control, two clones from the ancestral Degrader population and two clones from the ancestral Detoxifier population were picked and sequenced following the same workflow. All of the sequences can be accessed at the NCBI Sequence Read Archive (http://www.ncbi.nlm.nih.gov/sra) under Bioproject ID number PRJNA947372. The sequence alignment was performed between ancestors and evolved clones.

### Formulation of the mathematical model and the simulation protocols

The mathematical model was built to characterize the dynamics of a community composed of two strains (a Degrader and a Detoxifier). The model was built on seven assumptions based on our experimental system, and the first three are the key assumptions that we emphasized in the Results section.

1. The consortium may grow in two environments that enable two different types of interactions between the two strains. In the environment where the two strains engage in facultatively mutualistic cooperation (environment (Mutual.)), the Degrader strain grows on an initial substrate (S) and degrades S into a toxic intermediate (I) that is metabolized by the Detoxifier strain for detoxification. In the other environment, the two strains can grow independently (environment (Comp.)).
2. Autonomist may evolve from either population via horizontal gene transfer (HGT) in both environments. After an HGT event, the biomass of the evolved autonomous genotype follows a poison distribution with an expectation proportional to the biomass of both Degrader and Detoxifier^61,62^.
3. the evolved Autonomist population has a competitive edge (in terms of both the growth rate and carrying capacity) against the two cooperative specialists in the environment (Mutual.) but grows slower than them in the environment (Comp.).
4. the models were built on a well-mixed system.
5. the supply of naphthalene is sufficient in the environment (Mutual.), so the rate of S consumption remains constant, and the specific growth rate of Degrader remains constant.
6. the intracellular accumulation of I is negligible, so the consumption rate of I follows the first-order kinetics and the specific growth rate of Detoxifier is proportional to the concentration of I.
7. the degradation rate of S and the consumption rate of I are proportional to the biomass of the relevant population.

These assumptions allowed the building of minimal models that describe the growth dynamics of the consortium in the two environments. The population dynamics of one growth cycle in the environment (Mutual.) are given by

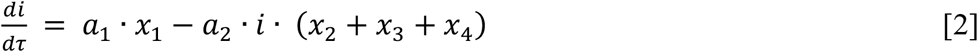

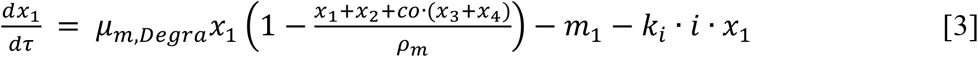

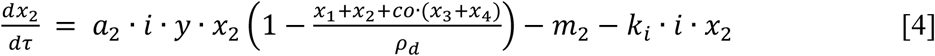

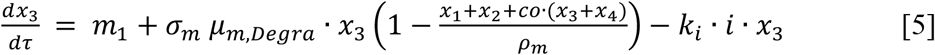

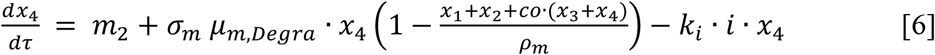

The population dynamics of one growth cycle in the environment (Comp.) are given by

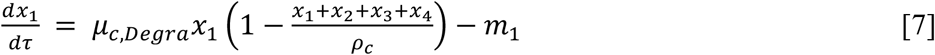

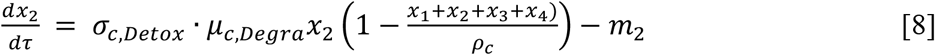

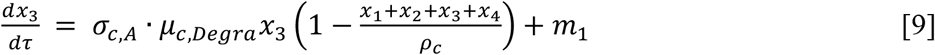

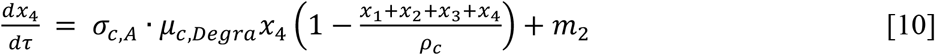

In the model, *τ* is the time. *x*_1_ is the biomass of Degrader while *x*_2_ is that of Detoxifier. *x*_3_ and *x*_4_ represent the biomass of Autonomist that evolve from Degrader and Detoxifier, respectively. *i* represents the concentration of the intermediate in the environment (Mutual.). *a*_1_ and *a*_2_ are the rates of the two reactions (S to I and I to P). *y* is the biomass yield coefficient of Detoxifier utilizing I. *μ*_*m,Degra*_ is the growth rate of Degrader in the environment (Mutual.). *σ*_*m*_ represents the relative growth rate of Autonomist to that of Degrader in the environment (Mutual.). Similarly, *μ*_*c,Degra*_ is the growth rate of Degrader in the environment (Comp.). *σ*_*c,Degra*_ and *σ*_*c,A*_ represent the relative growth rate of Detoxifier and Autonomist to that of Degrader in the environment (Comp.). *ρ*_*m*_ and *ρ*_*c*_ represent the carrying capacity of all populations in the environment (Mutual.) and (Comp.), respectively. *co* evaluates the competitive edge of Autonomist against Degrader and Detoxifier. *m*_1_ and *m*_2_ are the biomass of the evolved Autonomist after an HGT event, which follows a poison distribution with an expectation of *ε* ∙ *x*_1_*x*_2_, where *ε* evaluates the HGT rate.

To model the evolutionary dynamics of the mutualistic consortium, simulations were performed by serially passaging a consortium initially composed of Degrader and Detoxifier for 50 or 100 cycles with two conditions identical to our experiments: (1) the mutualism condition, in which the simulations were performed continuously using the equations [2] - [6] that characterize community dynamics in the environment (Mutual.); (2) the fluctuating condition, in which the simulations were performed alternately using the two equation systems. Since the evolved autonomous genotype can never dominate the community if only the environment (Comp.) is given according to the third assumption, the modeling of the competition condition was not performed. For each cycle, the consortium is allowed to grow to the early stationary phase (when the total biomass reaches 99.9% of the carrying capacity), mimicking the experiments. The consortium was then diluted to a novel cycle with a dilution factor of *d*.

To mathematically predict the experimental results, parameter values matching the experimental system^42^ (Table S3: the default values) were applied to the model. To investigate the key factors determining when fluctuating environments benefit the evolution of autonomy from mutualistic cooperation, 17501 parameter sets were generated by randomly picking the values of the 14 main parameters from the given ranges (Table S3). The ranges were chosen by incorporating at least ∼10-fold higher or lower than the default value. For each parameter set, simulations with the mutualism condition and the fluctuating condition were performed, and 20 replicates were set for each condition to capture the randomness in the evolution. The dynamics of different populations (i.e., *x*_1_, *x*_2_, *x*_3_, and *x*_4_) were recorded for further analysis. To identify the key factors determining the evolution of autonomy from mutualistic cooperation, several indices were defined for correlation analysis. First, *BCI*_*m*_ and *BCI*_*C*_ were defined to evaluate the cell density and coexistence level in the environment (Mutual.) and (Comp.), which are formulated as

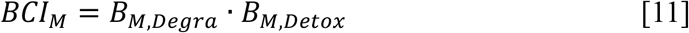

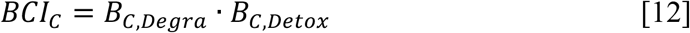

Here, *B*_*M,Degra*_ is the biomass of Degrader (*x*_1_) when the total biomass reaches 99.9% of the carrying capacity in the environment (Mutual.) while *B*_*M,Detox*_ is that of Detoxifier (*x*_2_). *B*_*C,Degra*_ and *B*_*C,Detox*_ represent the same values in the environment (Comp.). Thus, the two *BCI* values are proportional to the total biomass and community evenness (The closer the relative abundances of Degrader and Detoxifier, the higher evenness of the consortium, and the *BCI* values are also higher). Relative Biomass-Coexistence Index (*BCI*_*C/M*_) was then defined to characterize the relative cell density and coexistence levels in the environment (Comp.) than that in the environment (Mutual.), calculated by

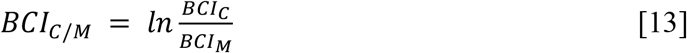

To evaluate the growth duration of the consortium in the two environments, *T*_*C*_ and *T*_*M*_ were introduced. *T*_*C*_ indicates the duration of the growth of the consortium to a total biomass reaching 99.9% of the carrying capacity in the environment (Comp.) while *T*_*M*_ indicates that duration in the environment (Mutual.). Accordingly, the Relative Duration of growing in the environment (Comp.) versus the environment (Mutual.) (*T*_*C/M*_) is defined as

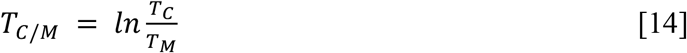

Finally, *P*_*fluc*_ and *P*_*mut*_ are used to quantify the probabilities of the evolution of autonomy in the fluctuating and mutualism conditions. *P*_*fluc*_ indicates the frequency of the simulations in which Autonomist dominates the community among 20 replicated simulations in the fluctuating condition while *P*_*mut*_ is that in the mutualism condition. The Fluctuation-Benefiting Index (*FBI*) quantifies whether environmental fluctuation is beneficial for the evolution of autonomy, calculated by

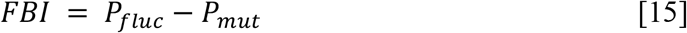

All simulations, as well as the downstream analysis, were performed using custom Mathematica scripts. The source codes used for the model analysis are publicly available (https://github.com/WMXgg/MDOLcode/tree/master/DOLEE%20model%20package).

### Quantification and statistical analyses

For the linear fit of the simulation data, data were first normalized by the maximum and minimum values, and the analyses were performed using the LinearModelFit function in *Wolfram Mathematica* (version 12.0) with default settings. The values of adjusted R-squared can be found in all related figures. Student’s T-test, Kolmogorov-Smirnov Test, and Pearson correlation analysis were calculated using the custom Mathematica scripts. To avoid the impact of the data size on significance analysis, the Student’s T-tests and Kolmogorov-Smirnov Tests of the simulation datasets were carried out with 1000 values randomly selected from each group sample. The statistical methods specific to each analysis are described in the corresponding figure legends.

### Replication and randomization

Replicate experiments have been performed for all key data shown in this study. Biological or technical replicate samples were randomized where appropriate. The numbers of replicates are listed in the related figure legends.

## Acknowledgments

We thank the members of the Wu lab at Peking University and Microbial Systems Ecology group at ETH Zürich for the critical discussions of this work. We thank Mr. Chunhui Hao from Oxford University for his helpful suggestions on the term usage in this study. We thank Dr. T. Juelich (UCAS, Beijing) for linguistic assistance during the preparation of this manuscript. We would like to express our gratitude to the organizers of the EMBO Workshop (Molecular Mechanisms in Evolution and Ecology) for providing us with the opportunity to present this work in the picturesque city of Heidelberg in the autumn of 2022. We also thank the Young Scholars Forum (2022) organized by the journal *mLife* for allowing us to showcase this research. We wish to extend our appreciation to the attendees of these conferences for their valuable feedback, which helped us a lot to improve our study. Miaoxiao Wang wishes to express his heartfelt gratitude to his wife, Yaxi Li, for her unwavering support and encouragement throughout this research project. This work was supported by National Key R&D Program of China (2018YFA0902100 and 2018YFA0902103), National Natural Science Foundation of China (91951204, 31761133006, 31770120, and 31770118), and Sino Swiss Science and Technology Cooperation (SSSTC) Program (IZLCZ0_206044).

## Author Contributions

All authors contributed intellectual input and assistance to this study. XC, MW, and YN conceived the study. XC and MW designed the evolution experiments. XC performed the experiments and analyzed the experimental data. MW constructed the ODE model, performed mathematical simulations, and analyzed the simulation data. MW drafted the manuscript with the help of XC. XC, YN, and XLW revised the manuscript. MW, YN, and XLW raised the funding for the project. All authors discussed the results and commented on the article.

