## supplemental file for "The evolution of autonomy from two cooperative specialists in fluctuating environments"

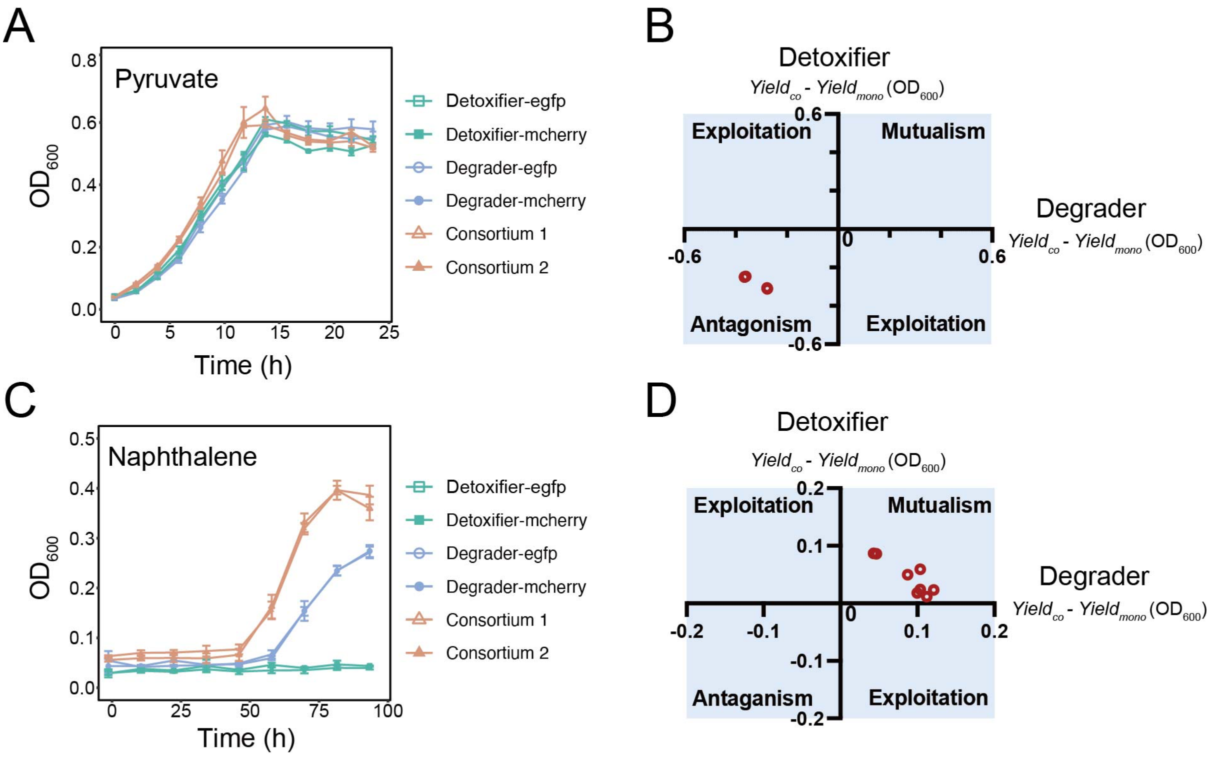


**Figure S1** Degrader and Detoxifier strains show antagonistic interaction upon co-culture using pyruvate (A-B), and mutualistic interaction upon co-culture using naphthalene (C-D). (A, C) The growth curve of Degrader and Detoxifier in monoculture and coculture, when pyruvate (A) or naphthalene (C) is used as the sole carbon source. Consortium 1 indicates the consortium composed of the Degrader strain labeled with mCherry and the Detoxifier strain labeled with eGFP, while Consortium 2 indicates the consortium composed of the Degrader strain labeled with eGFP and the Detoxifier strain labeled with mCherry. (B, D) Analysis of the interaction type based on the mono- and co-culture data. Interaction modes were defined by comparing the maximum cell density (quantified as optical density, $OD_{600}$) of monocultures and cocultures. (B) When pyruvate is used as the sole carbon source, the maximum cell densities of Degrader and Detoxifier in coculture were lower than that in the monoculture of both strains. Therefore, we propose that the two populations display antagonistic interaction. (D) When naphthalene is used as the sole carbon source, the maximum cell densities of Degrader and Detoxifier in coculture were higher than that in the monoculture of both strains. We thus propose that the two populations display mutualistic interaction.

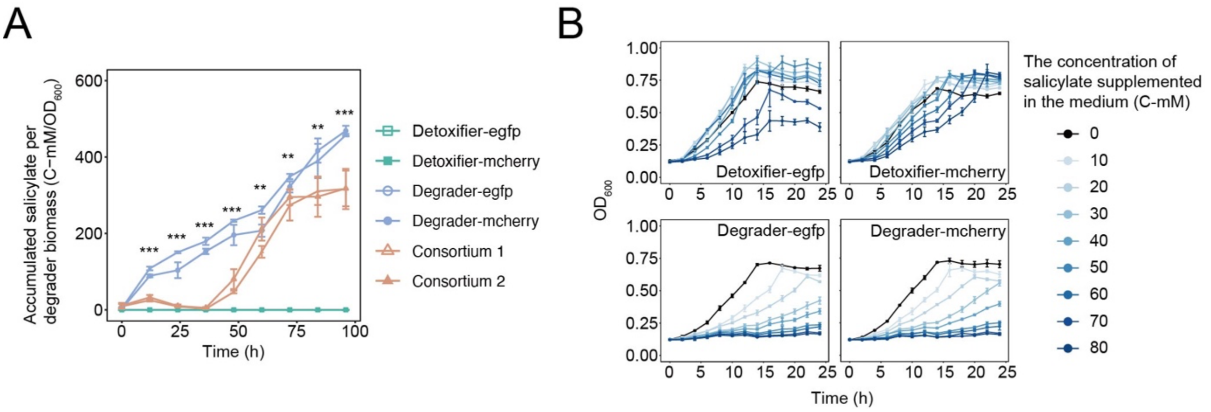


**Figure S2** Salicylate resistance and reduction ability of Degrader and Detoxifier. (A) Accumulated salicylate over time by degrading naphthalene in the minimal medium with naphthalene as the sole carbon source. Cocultures of Degrader and Detoxifier accumulate less salicylate in the medium than the monoculture of Degrader. (B) The growth curves of Degrader and Detoxifier by increasing salicylate concentrations in a minimal medium with 5 g/L pyruvate as the carbon source. The growth performance of Degrader and Detoxifier depends on initial salicylate concentrations added in the medium. The Detoxifier strains display higher salicylate resistance ability than the Degrader strains. The error bars represent the standard deviations for three biological replicates. The asterisks indicate significant differences (** $p < 0.01$; *** $p < 0.001$; Student's T-test, two-tailed).

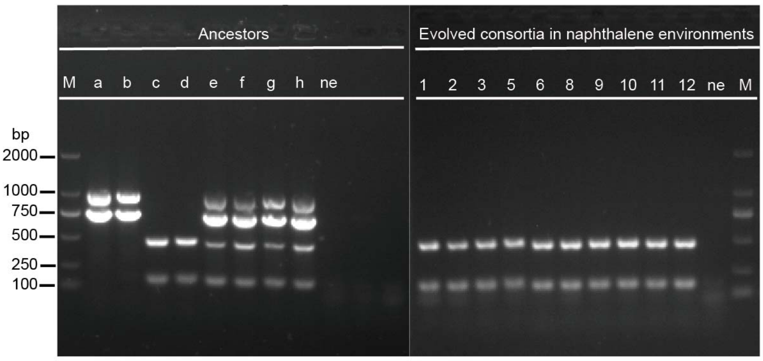


ne: $ddH_2O$
M: Marker
a: Ancestral Detoxifier-egfp
b: Ancestral Detoxifier-mcherry
c: Ancestral Degrader-egfp
d: Ancestral Degrader-mcherry
e: Ancestral Detoxifier-egfp + Ancestral Degrader-mcherry
f: Ancestral Detoxifier-mcherry + Ancestral Degrader-egfp
g: Ancestral Autonomist-egfp
h: Ancestral Autonomist-mcherry
1: Lin1
2: Lin2
3: Lin3
5: Lin5
6: Lin6
8: Lin8
9: Lin9
10: Lin10
11: Lin11
12: Lin12

**Figure S3** DNA electrophoresis shows that bands in all evolved lineages from the mutualism condition are consistent with the ancestral Degrader strains. The specific genes in Detoxifier, *nahG* and *nahTH*, were not detected in all lineages in the mutualism condition, demonstrating that Detoxifier is not detected in all lineages.

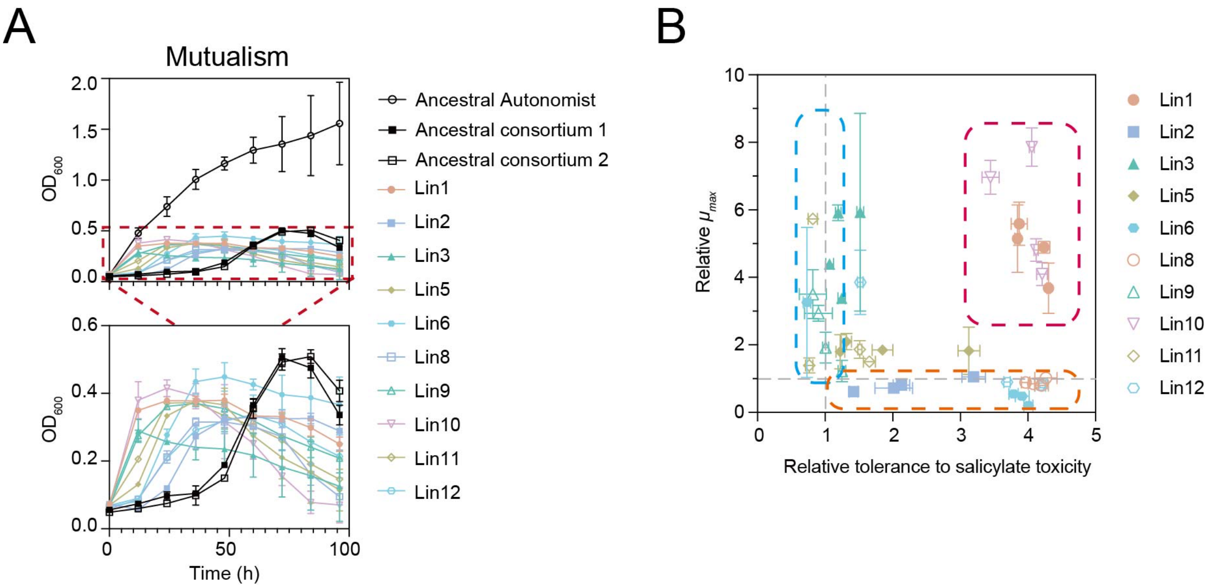


**Figure S4** The growth performance of the consortia and isolates evolved in the mutualism condition. (A) Growth kinetics of the consortia evolved in the mutualism condition using naphthalene as the sole carbon source. Ancestral consortium 1 indicates the consortium composed of the Degrader strain labeled with mCherry and the Detoxifier strain labeled with eGFP, while Ancestral consortium 2 indicates the consortium composed of the Degrader strain labeled with eGFP and the Detoxifier strain labeled with mCherry. (B) Growth rates and tolerance to salicylate toxicity of isolated clones from evolved consortia in the mutualism condition with naphthalene as the sole carbon source. The growth rate and tolerance to salicylate toxicity of the ancestral Degrader were measured as the control.

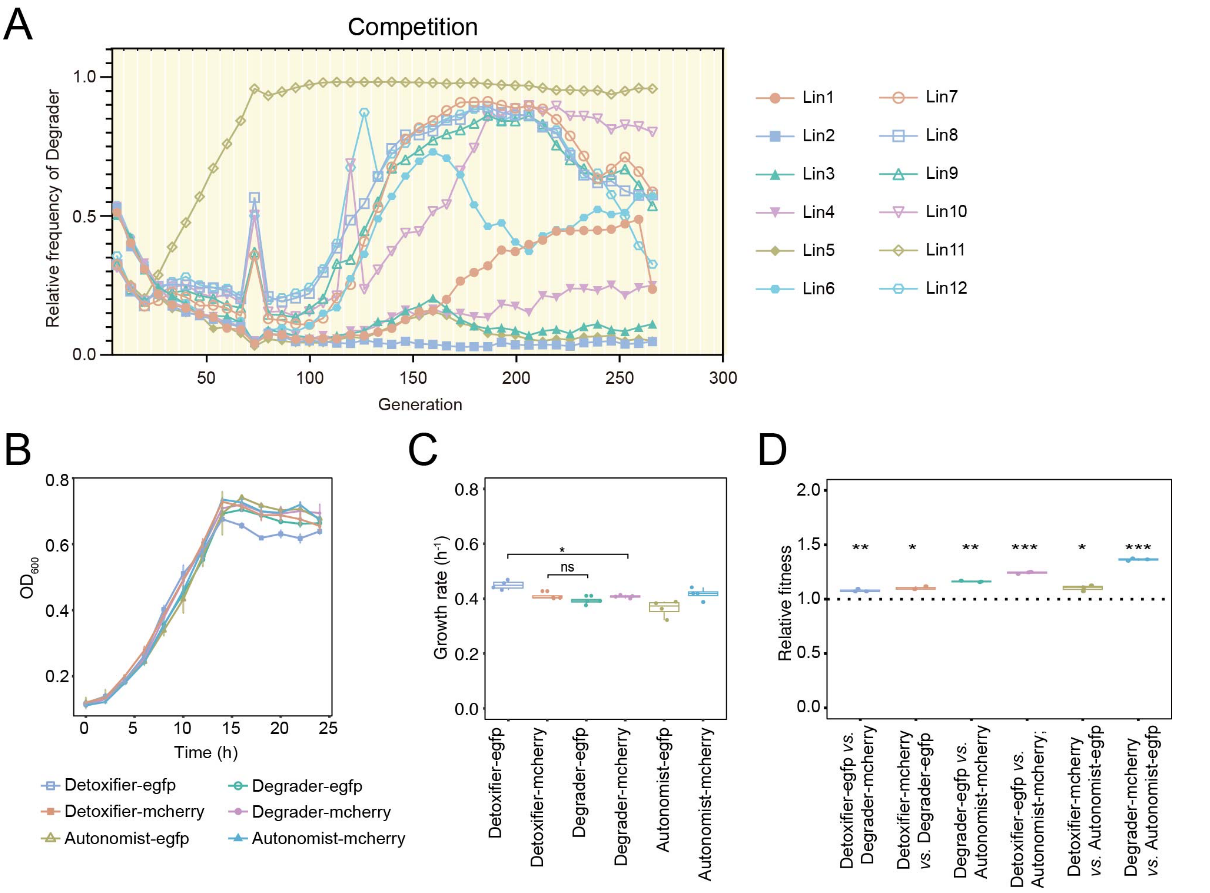


**Figure S5** The growth performance of ancestors and the dynamics of the mutualistic consortia in the evolution experiments in the minimal medium supplemented with 5 g/L pyruvate as the carbon source. (A) The dynamics of the community composition (denoted by the relative frequency of Degrader) in evolution experiments under the competition condition across all lineages. 12 independent replicates (i.e., lineages) are distinguished by the curve legends shown on the right of (A). (B) Growth kinetics and (C) growth rates of ancestors using pyruvate as the sole carbon source. (D) Competitive fitness between ancestral genotypes. $p$ values were derived from a Student's T-test. The asterisks indicate significant differences (ns $p > 0.05$, * $p < 0.05$, ** $p < 0.01$, *** $p < 0.001$).

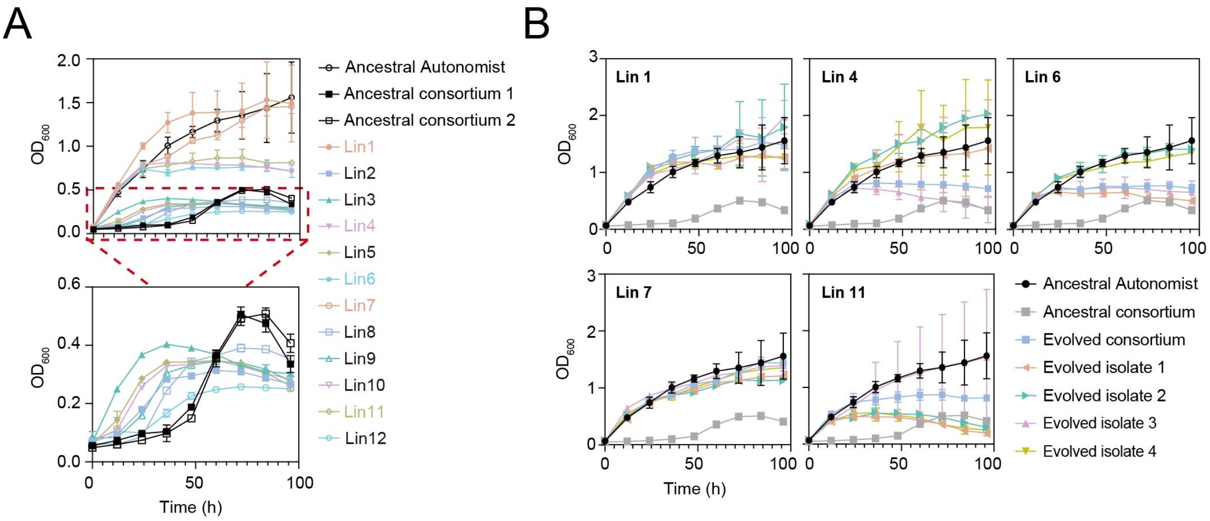


**Figure S6** The growth performance of the consortia and isolates evolved in the fluctuating condition. (A) Growth kinetics of the consortia evolved in the fluctuating condition using naphthalene as the sole carbon source. Ancestral consortium 1 indicates the consortium composed of the Degrader strain labeled with mCherry and the Detoxifier strain labeled with eGFP, while Ancestral consortium 2 indicates the consortium composed of the Degrader strain labeled with eGFP and the Detoxifier strain labeled with mCherry. The five lineages that have evolved under fluctuating conditions and exhibited higher maximum cell density ($OD_{max}$) than the ancestral consortia are highlighted in the caption on the right. (B) The growth curves of all the isolated clones in monoculture supplemented with naphthalene as the sole carbon source. The growth of the ancestral Autonomist strain and the ancestral mutualistic consortia were measured as controls. The error bars represent the standard deviations for three biological replicates.

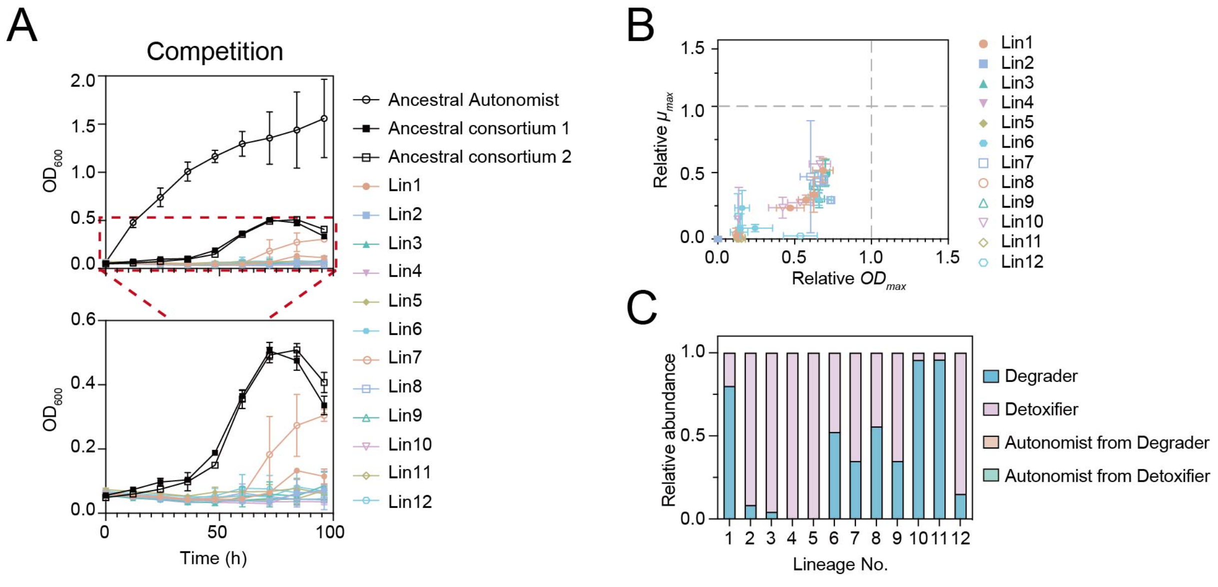


**Figure S7** The growth performance of the consortia and isolates evolved in the competition condition. (A) Growth kinetics of the consortia evolved in the competition condition using naphthalene as the sole carbon source. Ancestral consortium 1 indicates the consortium composed of the Degrader strain labeled with mCherry and the Detoxifier strain labeled with eGFP, while Ancestral consortium 2 indicates the consortium composed of the Degrader strain labeled with eGFP and the Detoxifier strain labeled with mCherry. (B) Comparison of the growth kinetics between the evolved isolates and ancestral consortia in the minimal medium with naphthalene as the sole carbon source. To determine their relative levels, we estimated the maximum cell density ($OD_{max}$) and the maximum growth rate ($\mu_{max}$) of the evolved isolates and those of the ancestral consortia ($n$ = 3). Dashed lines indicate levels of the ancestral consortia. (C) The relative frequencies of the different genotypes after the evolution in the competition condition, which demonstrates that Autonomist did not evolve in the competition condition.

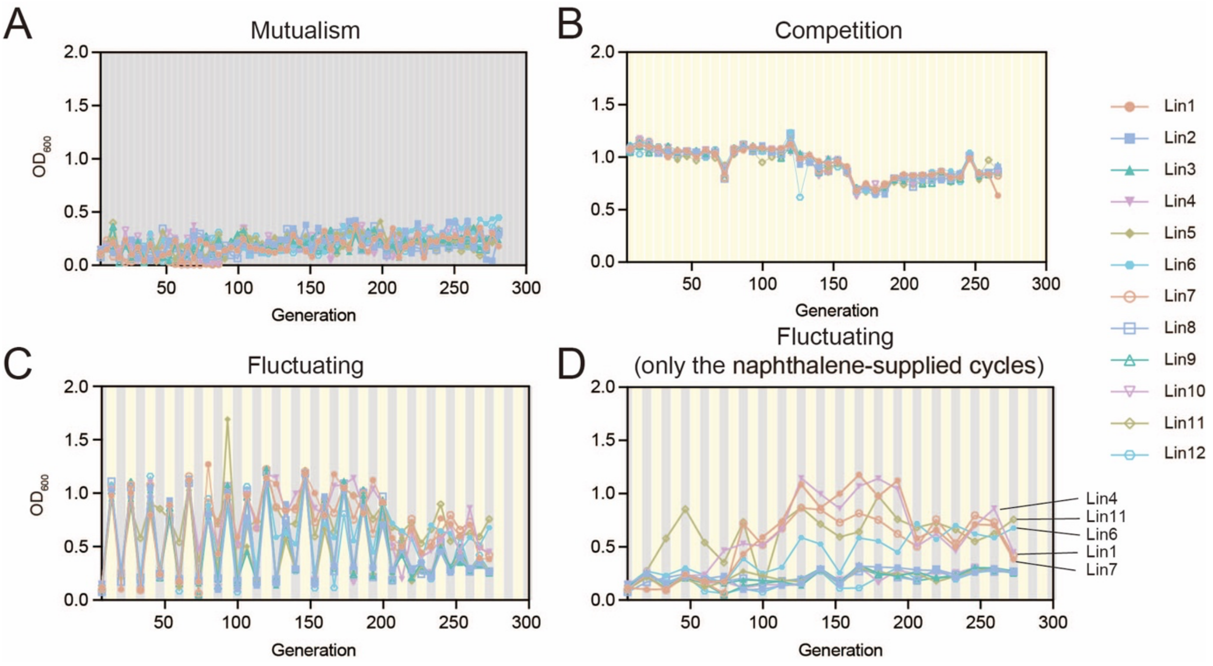


**Figure S8** Dynamics of cell density (quantified as optical density, $OD_{600}$) of the mutualistic consortia throughout the evolution experiment under the mutualism condition (A), the competition condition (B), and the fluctuating condition (C-D). The density of the consortia in the naphthalene-supplied cycles under the fluctuating condition is shown in (D), in which the five lineages in which Autonomist evolved under fluctuating conditions are labeled.

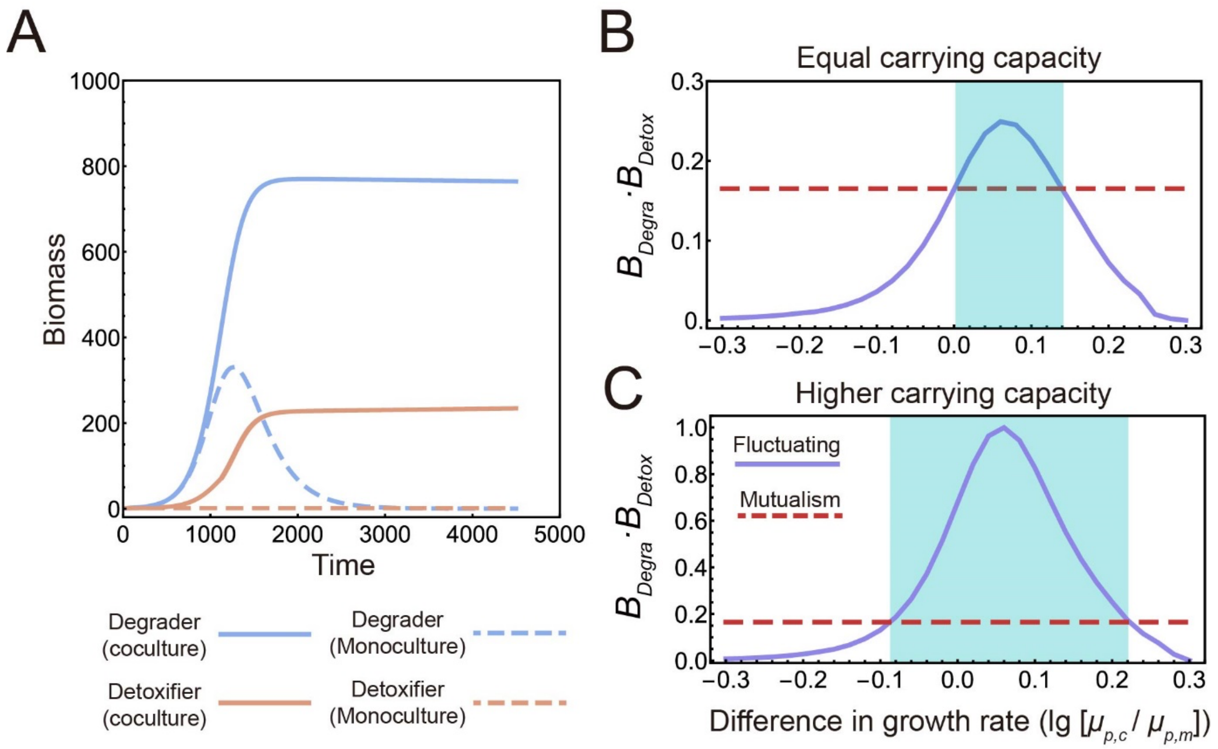


**Figure S9** Computational simulations using the parameters measured from our experiment systems (Table S3:default values). (A) Simulations of Degrader and Detoxifier in both monoculture and coculture revealed a mutualistic interaction between the two strains. In monoculture, Detoxifier was unable to grow alone; Degrader grew but experienced a decline after 1200 mins due to the accumulation and toxicity of the intermediate. In coculture, both strains exhibited faster growth and reached higher population densities compared to monoculture conditions. (B-C) The correlation between the relative growth rate of Detoxifier to that of Degrader in the environment (comp.) and the biomass-evenness level of the steady-state community was analyzed under two scenarios: (B) when the maximum carrying capacities of the two environments were the same; (C) the carrying capacity of the environment (Comp.) was set to be twice that of the environment (Mutual.). The biomass-evenness level ($B_{Degra} \cdot B_{Detox}$) was defined as the product of steady-state biomasses of Degrader and Detoxifier. This value was obtained from the replicates where Autonomist did not evolve and was proportional to the total biomass and community evenness (The closer the relative abundances of Degrader and Detoxifier, the higher evenness of the consortium, and the $BCI$ values are also higher).

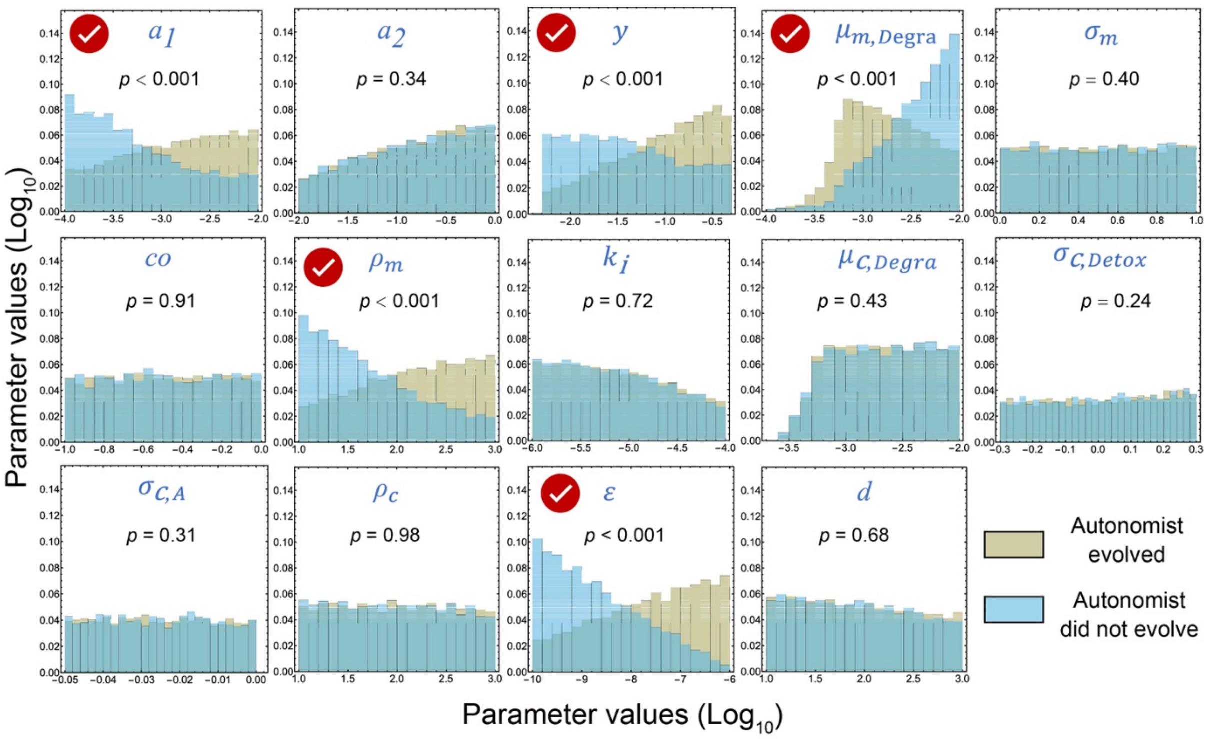


**Figure S10** Distributions of two groups of parameter values lead to two different types of simulation results under the mutualism condition: (1) The yellow histograms represent the parameter values of the 11636 sets resulting in the simulations in which Autonomist evolved during the simulations; (2) the blue histograms represent the parameter values of the 5865 sets resulting in the simulations in which Autonomist did not evolve during the simulations. The *p* values obtained from Kolmogorov-Smirnov Tests that compare the parameter values between the two groups are listed in each figure. Significant differences between the groups are indicated by markers placed on the corresponding parameters.

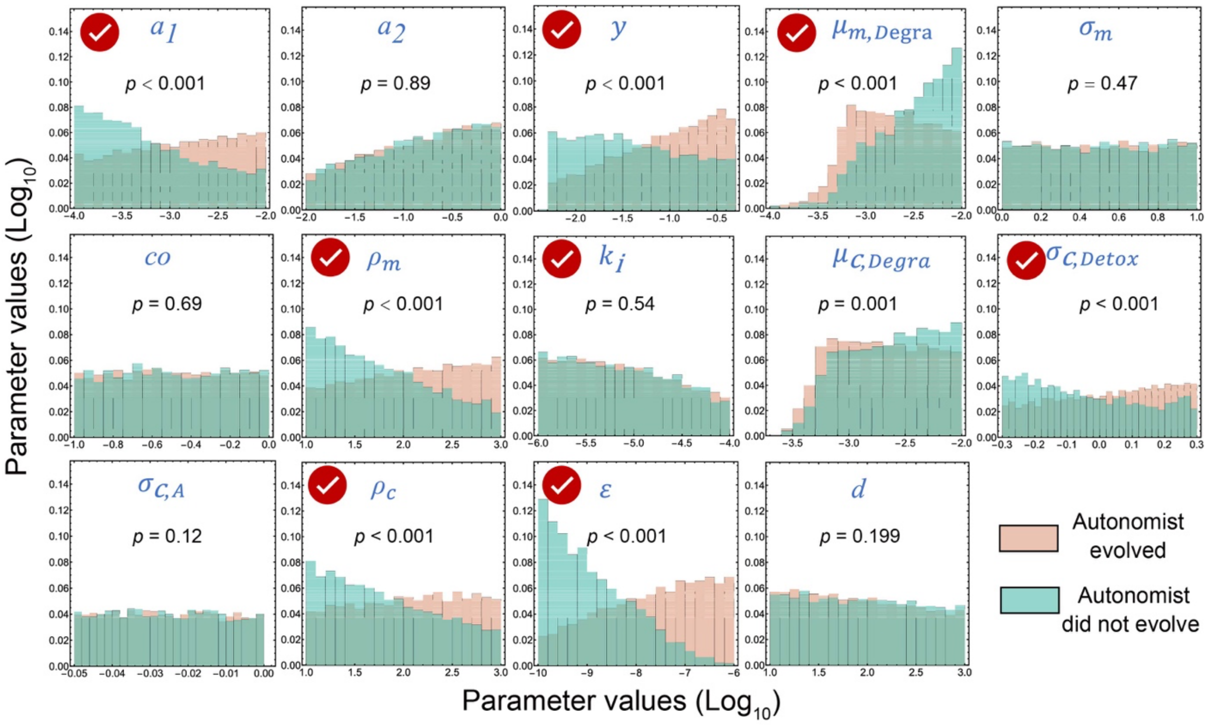


**Figure S11** Distributions of two groups of parameter values lead to two different types of simulation results under the fluctuating condition: (1) The pink histograms represent the parameter values of the 12902 sets resulting in the simulations in which Autonomist evolved during the simulations; (2) the green histograms represent the parameter values of the 4599 sets resulting in the simulations in which Autonomist did not evolve during the simulations. The *p*-values obtained from Kolmogorov-Smirnov Tests that compare the parameter values between the two groups are listed in each figure. Significant differences between the groups are indicated by markers placed on the corresponding parameters.

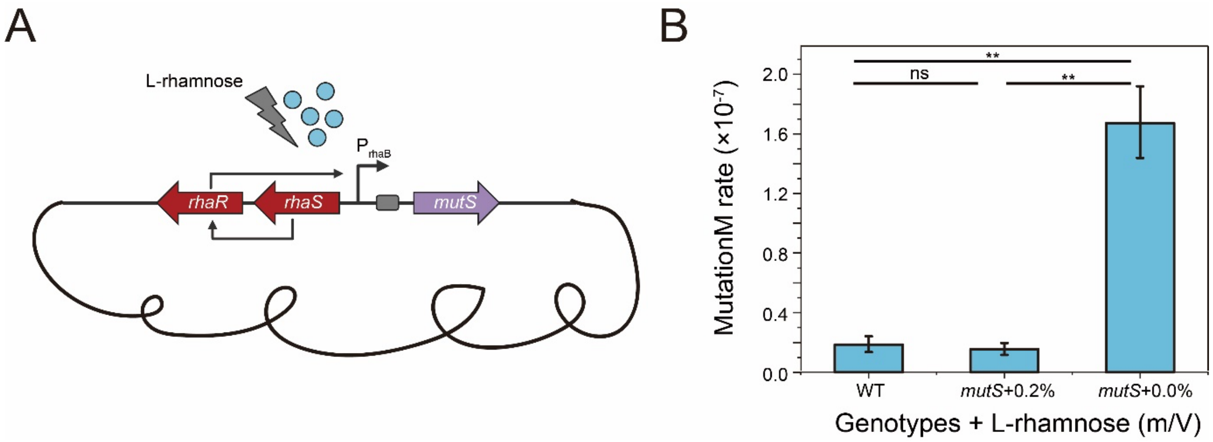


**Figure S12** Inducible expression of *mutS* in our engineered strains. (A) The L-rhamnose inducible promoter *rhaSR-PrhaBAD* was inserted in front of the *mutS* to control the expression of *mutS*. (B) The mutation rates of the engineered strains without this modification (WT), as well as the modified strains in the presence and absence of the L-rhamnose. The mutation rates were measured by Luria and Delbruck fluctuation analysis. We observed that the mutation rate in the strain lacking the expression of *mutS* was significantly higher than that in the strain expressing *mutS,* as well as significantly higher than that in WT (ns $p > 0.05$; ** $p < 0.0001$).

**Table S1** Mapping the barcode sequences of evolved strains under the mutualism condition onto the barcodes of the ancestral strains.

| Lineage No. | Clones No. | Match Degrader's barcode? Degrader's barcode: ACCTACCTTAATGGTTGTTGGTCTTAGG | Match Detoxifier's barcode? Detoxifier's barcode: ACTTACGGGAATTTAGGTGGAACTGCAG | Match Autonomist's barcode? Autonomist's barcode: CCCGACTTGGATGTGGGTATATCTGAAT |
|---|---|---|---|---|
| Lin1 | 1 | Yes | No | No |
| | 2 | Yes | No | No |
| | 3 | Yes | No | No |
| | 4 | Yes | No | No |
| Lin2 | 1 | Yes | No | No |
| | 2 | Yes | No | No |
| | 3 | Yes | No | No |
| | 4 | Yes | No | No |
| Lin3 | 1 | Yes | No | No |
| | 2 | Yes | No | No |
| | 3 | Yes | No | No |
| | 4 | Yes | No | No |
| Lin5 | 1 | Yes | No | No |
| | 2 | Yes | No | No |
| | 3 | Yes | No | No |
| | 4 | Yes | No | No |

| | | | | |
|---|---|---|---|---|
| Lin6 | 1 | Yes | No | No |
| | 2 | Yes | No | No |
| | 3 | Yes | No | No |
| | 4 | Yes | No | No |
| Lin8 | 1 | Yes | No | No |
| | 2 | Yes | No | No |
| | 3 | Yes | No | No |
| | 4 | Yes | No | No |
| Lin9 | 1 | Yes | No | No |
| | 2 | Yes | No | No |
| | 3 | Yes | No | No |
| | 4 | Yes | No | No |
| Lin10 | 1 | Yes | No | No |
| | 2 | Yes | No | No |
| | 3 | Yes | No | No |
| | 4 | Yes | No | No |
| Lin11 | 1 | Yes | No | No |
| | 2 | Yes | No | No |
| | 3 | Yes | No | No |
| | 4 | Yes | No | No |
| Lin12 | 1 | Yes | No | No |

| | | | |
|---|---|---|---|
| 2 | Yes | No | No |
| 3 | Yes | No | No |
| 4 | Yes | No | No |

**Table S2** Mapping the barcode sequences of evolved Autonomist under the fluctuating condition onto the barcodes of the ancestral strains.

| Lineage No. | Clones No. | Match Degrader's barcode? Degrader's barcode: ACCTACCTTAATGGTTGTTGGTCTTAGG | Match Detoxifier's barcode? Detoxifier's barcode: ACTTACGGGAATTTAGGTGGAACTGCAG | Match Autonomist's barcode? Autonomist's barcode: CCCGACTTGGATGTGGGTATATCTGAAT |
|---|---|---|---|---|
| Lin1 | 1 | Yes | No | No |
| | 2 | Yes | No | No |
| | 3 | Yes | No | No |
| | 4 | Yes | No | No |
| Lin4 | 1 | Yes | No | No |
| | 2 | Yes | No | No |
| | 3 | Yes | No | No |
| | 4 | Yes | No | No |
| Lin6 | 1 | Yes | No | No |
| | 2 | Yes | No | No |
| | 3 | Yes | No | No |
| | 4 | Yes | No | No |
| Lin7 | 1 | No | Yes | No |
| | 2 | No | Yes | No |
| | 3 | No | Yes | No |
| | 4 | No | Yes | No |

| | | | | |
|---|---|---|---|---|
| Lin11 | 1 | No | Yes | No |
| | 2 | No | Yes | No |
| | 3 | No | Yes | No |
| | 4 | No | Yes | No |

**Table S3** Variables and parameters used in the model.

| Parameters | Definitions | Unit | Default values | Search Range | Sources |
|---|---|---|---|---|---|
| **Variables** | | | | | |
| $x_1$ | The biomass of Degrader | $10^5$ cells$^{-1}$ | N.A. | N.A. | N.A. |
| $x_2$ | The biomass of Detoxifier | $10^5$ cells$^{-1}$ | N.A. | N.A. | N.A. |
| $x_3$ | The biomass of Autonomist evolved from Degrader | $10^5$ cells$^{-1}$ | N.A. | N.A. | N.A. |
| $x_4$ | The biomass of Autonomist evolved from Detoxifier | $10^5$ cells$^{-1}$ | N.A. | N.A. | N.A. |
| $i$ | The concentration of the intermediate in the environment (Mutual.) | mol | N.A. | N.A. | N.A. |
| $\tau$ | Simulation time | min$^{-1}$ | N.A. | N.A. | N.A. |
| **Parameters regarding the dynamics in the environment (Mutual.)** | | | | | |
| $a_1$ | Rate of the first reaction | mol·min$^{-1}$·$10^5$ cells$^{-1}$ | 0.005 | $10^{-4}$ – $10^{-2}$ | Experiments |
| $a_2$ | Specific rate of the second reaction | min$^{-1}$·$10^5$ cells$^{-1}$ | 0.02 | $10^{-2}$ – $10^0$ | Experiments |
| $y$ | Yield coefficient of Detoxifier | $10^5$ cells · mol | 0.1 | $10^{-2}$ – $10^0$ | Experiments |
| $\mu_{m,Degra}$ | Growth rate of Degrader in the mutualistic system | min$^{-1}$ | 0.003 | $10^{-4}$ – $10^{-2}$ | Experiments |
| $\sigma_m$ | Relative growth rate of Autonomist to that of Degrader in the environment (Mutual.) | N.A. | 1.7 | $10^0$ – $10^1$ | Experiments |

| | | | | | |
|---|---|---|---|---|---|
| $co$ | Competitive edge of Autonomist in the environment (Mutual.) | N.A. | 1/3 | $10^{-1}$ – $10^{0}$ | Experiments |
| $k_i$ | Toxic coefficient of the intermediate | $min^{-1}\cdot mol\cdot 10^{5}$ $cells^{-1}$ | 0.0000016 | $10^{-7}$ – $10^{-5}$ | Experiments |
| $\rho_m$ | Maximum carrying capacity in the environment (Mutual.) | $10^{5}$ cells | 1000 | $10^{1}$ – $10^{4}$ | Experiments |
| **Parameters regarding the dynamics in the environment (Comp.)** | | | | | |
| $\mu_{c,Degra}$ | Growth rate of Degrader in the competition system | $min^{-1}$ | 0.005 | $5\times10^{-4}$ – $5\times10^{-2}$ | Experiments |
| $\sigma_{c,Detox}$ | Relative growth rate of Detoxifier to that of Degrader in the environment (Comp.) | N.A. | 1 | 0.5 – 2 | Experiments |
| $\sigma_{c,A}$ | Relative growth rate of Autonomist to that of Degrader in the environment (Comp.) | N.A. | 1 | $10^{-1}$ – $10^{0}$ | Experiments |
| $\rho_c$ | Maximum carrying capacity in the environment (Comp.) | $10^{5}$ cells | 2000 | $10^{1}$ – $10^{4}$ | Experiments |
| **Other Parameters** | | | | | |
| ε | Specific rate of horizontal gene transfer | $10^{10}$ $cells^{-1}$ | $5\times10^{-11}$ | $10^{-12}$ – $10^{-8}$ | [1] |
| $d$ | Dilution factor | N.A. | $10^{3}$ | $10^{1}$ – $10^{5}$ | Experiments |
| $Cn$ | The number of cycles | N.A. | 20 | 100 | Experiments |
| **Initial conditions** | | | | | |
| $[x_{1,0}, x_{2,0}, x_{3,0}, x_{4,0}]$ | The initial density of the cells | $10^{5}$ cells | [1, 1, 0, 0] | N.A. | Experiments |

| | | | | | |
|---|---|---|---|---|---|
| $i_0$ | The initial concentration of the intermediate | mol | 0 | N.A. | N.A. |

**Table S4** Strains and plasmids used in this study

| Strains or plasmids | Features | Source |
|---|---|---|
| **Strains** | | |
| *E.coli* Trans5α | $F^-$ φ80 *lac ZΔM15 Δ(lacZYA-arg F) U169 endA1 recA1 hsdR17*($r_k^-$, $m_k^+$) *supE44λ⁻ thi -1 gyrA96 relA1 phoA* | TransGen |
| *E.coli* HB101 | $F^-$ *mcrB mrr hsdS20*(rB mB) *recA13 leuB6 ara-14 proA2 lacY1 galK2 xyl-5 mtl-1 rpsL20*(Smr) *glnV44* | Lab of Professor Lin Min, CAAS |
| *P. stutzeri* AN1111 | *P. stutzeri* AN10△Pnah::Ptac△Psal::Ptac△nahW△catA△nahR::LacI | [2] |
| *P. stutzeri* AN1100 | AN1111△*nahG::A*△*nahTH::C* | [2] |
| *P. stutzeri* AN0011 | AN1111△*nahA::A*△*nahC::C* | [2] |
| *P. stutzeri* AN1111-barcode16-mCherry | The ancestral Autonomist strain; *P. stutzeri* AN1100 containning pJMbarcode16-mCherry | This study |
| *P. stutzeri* AN1111-barcode16-eGFP | The ancestral Autonomist strain; *P. stutzeri* AN1100 containning pJMbarcode16-eGFP | This study |
| *P. stutzeri* AN1100-barcode9-mCherry | *P. stutzeri* AN1100 containning pJMbarcode9-mCherry | This study |
| *P. stutzeri* AN1100-barcode9-eGFP | *P. stutzeri* AN1100 containning pJMbarcode9-eGFP | This study |
| *P. stutzeri* AN0011-barcode8-mCherry | *P. stutzeri* AN1100 containning pJMbarcode8-mCherry | This study |
| *P. stutzeri* AN0011-barcode8-eGFP | *P. stutzeri* AN1100 containning pJMbarcode8-eGFP | This study |
| *P. stutzeri* AN1100-barcode9-mCherry-mutS | The ancestral Degrader strain; *P. stutzeri* AN1100-barcode9-mCherry△*mutS::rhaSR-PrhaBAD-mutS* | This study |
| *P. stutzeri* AN1100-barcode9-eGFP-mutS | The ancestral Degrader strain; *P. stutzeri* AN1100-barcode9-eGFP△*mutS::rhaSR-PrhaBAD-mutS* | This study |
| *P. stutzeri* AN0011-barcode8-mCherry-mutS | The ancestral Detoxifier strain; *P. stutzeri* AN0011-barcode8-mCherry△*mutS::rhaSR-PrhaBAD-mutS* | This study |

| | | |
|---|---|---|
| *P. stutzeri* AN0011-barcode8-eGFP-mutS | The ancestral Detoxifier strain; *P. stutzeri* AN0011-barcode8-eGFP$\Delta$*mutS::rhaSR-PrhaBAD-mutS* | This study |
| **Plasmids** | | |
| pRK2013 | Helper plasmid for conjugation; $Km^R$ | Lab of Professor Lin Min, CAAS[3] |
| pTNS3 | Helper plasmid; $Amp^R$ | Addgene[4] |
| pJM220 | pUC18T-miniTn7T-gm-rhaSR-PrhaBAD; $Gm^R$ | Addgene[4] |
| pJMbarcode8-mCherry | *barcode8-mCherry* on pJM220 | This study |
| pJMbarcode8-eGFP | *barcode8-eGFP* on pJM220 | This study |
| pJMbarcode9-mCherry | *barcode9-mCherry* on pJM220 | This study |
| pJMbarcode9-eGFP | *barcode9-eGFP* on pJM220 | This study |
| pJMbarcode16-mCherry | *barcode16-mCherry* on pJM220 | This study |
| pJMbarcode16-eGFP | *barcode16-eGFP* on pJM220 | This study |
| pK18mobsacB-*rhaSR-PrhaBAD-mutS* | *rhaSR-PrhaBAD-mutS* on pK18mobsacB | This study |

**Table S5** Primers used in this study

| Primers | Oligonucleotides (5'→3') | Description |
|---|---|---|
| nahA-1 | catcgacttgattgccgcgtcgt | Detecting the presence or absence of the *nahA* gene in evolved consortia and isolates |
| nahA-2 | tggatccgcaaaactggggcaatagtcc | Detecting the presence or absence of the *nahA* gene in evolved consortia and isolates |
| nahC-1 | ggctcaatgagccagccggaag | Detecting the presence or absence of the *nahC* gene in evolved consortia and isolates |
| nahC-2 | cccgaccgagatattctggggcc | Detecting the presence or absence of the *nahC* gene in evolved consortia and isolates |
| nahG-1 | gagtcgccagtatttcccccaccagc | Detecting the presence or absence of the *nahG* gene in evolved consortia and isolates |
| nahG-2 | cagagtaccgctgcgatcttctaatcggc | Detecting the presence or absence of the *nahG* gene in evolved consortia and isolates |
| nahTH-1 | ggtcgtgatagaagatcgccttgcccag | Detecting the presence or absence of the *nahTH* gene in evolved consortia and isolates |
| nahTH-2 | gaatgcgggcgtatgagctgcaag | Detecting the presence or absence of the *nahTH* gene in evolved consortia and isolates |
| Pc_R | ttctcgaggaattcctgcagtgccgatacaagaacaacagcg | Amplifying the *eGFP* gene |
| eGFP_F | tcactagtggatcccccgggTTACTTGTACAGTTCGTCCATGCC | Amplifying the *eGFP* gene |
| morange-F | tcactagtggatcccccgggTTACTTGTACAGTTCGTCCATGCCG | Amplifying the *mCherry* gene |
| Pc_R | ttctcgaggaattcctgcagtgccgatacaagaacaacagcg | Amplifying the *mCherry* gene |
| p18-minitn7-F | ctgcaggaattcctcgagaagc | Amplifying the backbone of pJM220 with barcode |
| p18-minitn7-R | cccgggggatccactagtg | Amplifying the backbone of pJM220 with |

| | | |
|---|---|---|
| | | barcode |
| junP-F | TTACGAACCGAACAGGCTTATG | Constructing pJMbarcode-fluorescence |
| junP-R | GGCAGGATAGGTGAAGTAGGC | Constructing pJMbarcode-fluorescence |
| pglmS-up | atgccgatcaatgtcggact | Verifying the knock-in result |
| pglmS-down | atcgtctttgccgatcgtgaag | Verifying the knock-in result |
| cexu-R | cgttgctgctccataacatcaaac | Verifying the knock-in result |
